# A frameshift in *Yersinia pestis rcsD* leads to expression of a small HPt variant that alters canonical Rcs signalling to preserve flea-mammal plague transmission cycles

**DOI:** 10.1101/2022.10.24.512971

**Authors:** Xiao-Peng Guo, Hai-Qin Yan, Wenhui Yang, Zhe Yin, Viveka Vadyvaloo, Dongsheng Zhou, Yi-Cheng Sun

**Affiliations:** MOH key laboratory of Systems Biology of Pathogens, Institute of Pathogen Biology, Chinese Academy of Medical Sciences and Peking Union Medical College, 9, Dongdan Santiao, Dongcheng District, Beijing, 100730, China; Department of Basic Medical Sciences, Anhui Key Laboratory of Infection and Immunity, Bengbu Medical College, Bengbu, 233030, China; State Key Laboratory of Pathogen and Biosecurity, Beijing Institute of Microbiology and Epidemiology, 100071, Beijing, China; Paul G. Allen School for Global Health, Washington State University, Pullman, WA 99164, USA

## Abstract

Multiple genetic changes in the enteric pathogen *Yersinia pseudotuberculosis* have driven the emergence of *Yesinia pestis*, the arthropod-borne, etiological agent of plague. These include developing the capacity for biofilm-dependent blockage of the flea foregut to enable transmission by flea bite. Previously, we showed that pseudogenisation of *rcsA*, encoding a component of the Rcs signalling pathway, is an important evolutionary step facilitating *Y. pestis* flea-borne transmission. Additionally, *rcsD*, another important gene in the Rcs system, harbours a frameshift mutation. Here, we demonstrated that this *rcsD* mutation resulted in predominant production of a small protein composing the C-terminal RcsD histidine-phosphotransferase domain (designated RcsD-Hpt) and low levels of full-length RcsD. Genetic analysis revealed that the *rcsD* frameshift mutation followed the emergence of *rcsA* pseudogenisation. It further altered the canonical Rcs phosphorylation signal cascade, fine-tuning biofilm production to be conducive with retention of the *pgm* locus in modern lineages of *Y. pestis*. Taken together, our findings suggest that a frameshift mutation in *rcsD*, is an important evolutionary step that fine-tuned biofilm production to ensure perpetuation of flea-mammal plague transmission cycles.

## Introduction

Approximately 6,000–7,000 years ago, *Yersinia pestis* evolved to be an arthropod-borne pathogen from its ancestor *Yersinia pseudotuberculosis* (Cui et al., 2013; Rasmussen et al., 2015; Spyrou et al., 2018). Despite their recent divergence, these species have markedly different life cycles. *Y. pseudotuberculosis* is transmitted by the faecal-oral route and usually causes a mild, self-limiting enteric disease in mammals (Putzker et al., 2001). *Y. pestis*, uniquely amongst enteric Gram-negative bacteria, is highly virulent and relies on flea-borne transmission (Hinnebusch, 2005). The co-occurrence of both ancestor and descendant *Yersinia* species provides an exemplary model to study the evolution of bacterial pathogens (Hinchliffe et al., 2003; Hinnebusch, 1997; Wren, 2003).

*Y. pestis* is transmitted by flea bites via a crude regurgitation mechanism (Hinnebusch and Erickson, 2008). After entering the flea gut, the planktonic *Y. pestis* form large aggregates and colonise the proventriculus. When infected fleas feed again as early as 1-3 days post infection, the bacterial mass in the proventriculus is regurgitated into the bite site, leading to a phenomenon referred to as early phase transmission (Bosio et al., 2020; Dewitte et al., 2020; Eisen et al., 2007; Hinnebusch and Erickson, 2008). Concurrently, *Y. pestis* continues to multiply in the flea digestive tract and forms HmsHFRS-dependent biofilms in the proventriculus, which blocks flea feeding (Abu Khweek et al., 2010). Continuous attempts to feed by the starved flea promotes reflux of bacteria-contaminated blood to the bite site, a phenomena termed biofilm-dependent transmission (Bland et al., 2018). In contrast to the low efficiency of early phase transmission, which usually requires several infected fleas simultaneously feeding on a naïve host, biofilm-dependent late-stage transmission is highly efficient, and a single blocked flea has high potential for transmission (Bosio et al., 2020).

*Y. pestis* diverged from *Y. pseudotuberculosis* through a series of gene gains, gene losses and genome rearrangements (Cao et al., 2022; Hinchliffe et al., 2003; McNally et al., 2016; Rascovan et al., 2019; Sun et al., 2014). Acquisition of the *ymt* gene enabled *Y. pestis* to survive and colonise the flea midgut to sustain flea-borne plague through expansion of its mammalian host range (Bland et al., 2021; Sun et al., 2014). Loss of the three genes, *rcsA, pde2* and *pde3*, altered the c-di-GMP signalling pathway, which increased biofilm-forming capability (Sun et al., 2008; Sun et al., 2014). These four genetic changes enabled the *Y. pseudotuberculosis* progenitor strain to form biofilms in the proventriculus of fleas, promoting a flea-borne transmission modality (Sun et al., 2014).

The Rcs phosphorelay system, a non-orthodox two-component signal transduction system, consists of a hybrid sensor kinase RcsC, the phosphotransfer protein RcsD, and a response regulator RcsB (Guo and Sun, 2017; Wall et al., 2018). In *Enterobacteriaceae*, the outer membrane lipoprotein, RcsF, senses outer membrane (OM)- and peptidoglycan-related stress (Smith et al., 2021; Tata et al., 2021). The inner membrane protein, IgaA, subsequently relays these signals to RcsD, which activates the Rcs phosphorelay system (Cho et al., 2014; Wall et al., 2020). Autophosphorylated RcsC transfers a phosphate group to a conserved histidine residue in the C-terminal histidine-phosphotransferase (HPt) domain of RcsD, which is finally transferred to RcsB (Takeda et al., 2001). Phosphorylated RcsB acts either alone or in combination with auxiliary proteins to regulate expression of target genes (Clarke, 2010; Huesa et al., 2021; Pannen et al., 2016). Functional RcsA works in concert with RcsB as a heterodimer to inhibit *Y. pseudotuberculosis* biofilm formation, in part by repressing expression of the c-di-GMP synthesis genes *hmsT* and *hmsD* (Bobrov et al., 2011; Fang et al., 2015; Guo et al., 2015; Sun et al., 2012; Sun et al., 2011). In *Y. pestis*, however, RcsA (RcsA_pe_) was disrupted by acquiring a 30 bp repeat insertion sequence, leading to enhanced capacity for biofilm formation (Sun et al., 2008). *rcsD* in *Y. pestis* (*rcsD*_pe_) is a putative pseudogene due to a 1 bp deletion, but it retains a limited ability to modulate biofilm formation *in vitro* (Sun et al., 2008).

Here, we investigated the functional consequences of the frameshift mutation in *rcsD*_pe_ and the mechanism by which it modulates activity of the Rcs signalling system. We found that the *rcsD*_pe_ variant produces a 103-amino acid protein containing the C- terminal HPt domain of RcsD, designated as RcsD-Hpt, and that this protein plays a dominant role in Rcs signalling in *Y. pestis*. Frameshifted *rcsD* alters Rcs signal transduction, subsequently modulating the expression of dozens of genes and the capacity for biofilm formation in *Y. pestis*. These evolutionary events may represent an important step in the emergence of ubiquitous branches of *Y. pestis* that can capably maintain plague outbreaks.

## Results

### *rcsD*_pe_ negatively regulates, while *rcsD*_pstb_ positively regulates, biofilm formation in *Y. pestis*

RcsD, an inner membrane protein, has an HPt domain at its C-terminus (Takeda et al., 2001; Wall et al., 2018). The *rcsD* gene in *Y. pestis* has undergone a frameshift after codon 642 due to single nucleotide deletion (Figure 1a). However, *rcsD*_pe_ is still functional as deletion of the N-terminal region (1–1846 bp, Δ*rcsD*_N-term_) of *rcsD*_pe_ reduced biofilm formation and Congo red (CR) pigmentation (Figure 1b and 1c) (Sun et al., 2008), while replacement of *rcsD*_pe_ with *rcsD*_pstb_ significantly increased biofilm formation (Figure 1b) (Sun et al., 2008). To further characterise the differences between *rcsD*_pe_ and *rcsD*_pstb_, we expressed high levels of each gene in a *Y. pestis* wild type and *rcsD* N-terminal deletion strain, respectively (Figure 1b and 1c). Consistent with our previous report (Sun et al., 2008), overexpression of *rcsD*_pstb_ significantly increased CR pigmentation and biofilm formation *in vitro* (Figure 1b and 1c). By contrast, overexpression of *rcsD*_pe_ decreased the CR phenotype and formation of biofilm (Figure 1b and 1c), indicating that *rcsD*_pe_ might play a different role to *rcsD*_pstb_ in the modulation of Rcs signalling.

**Figure 1.**
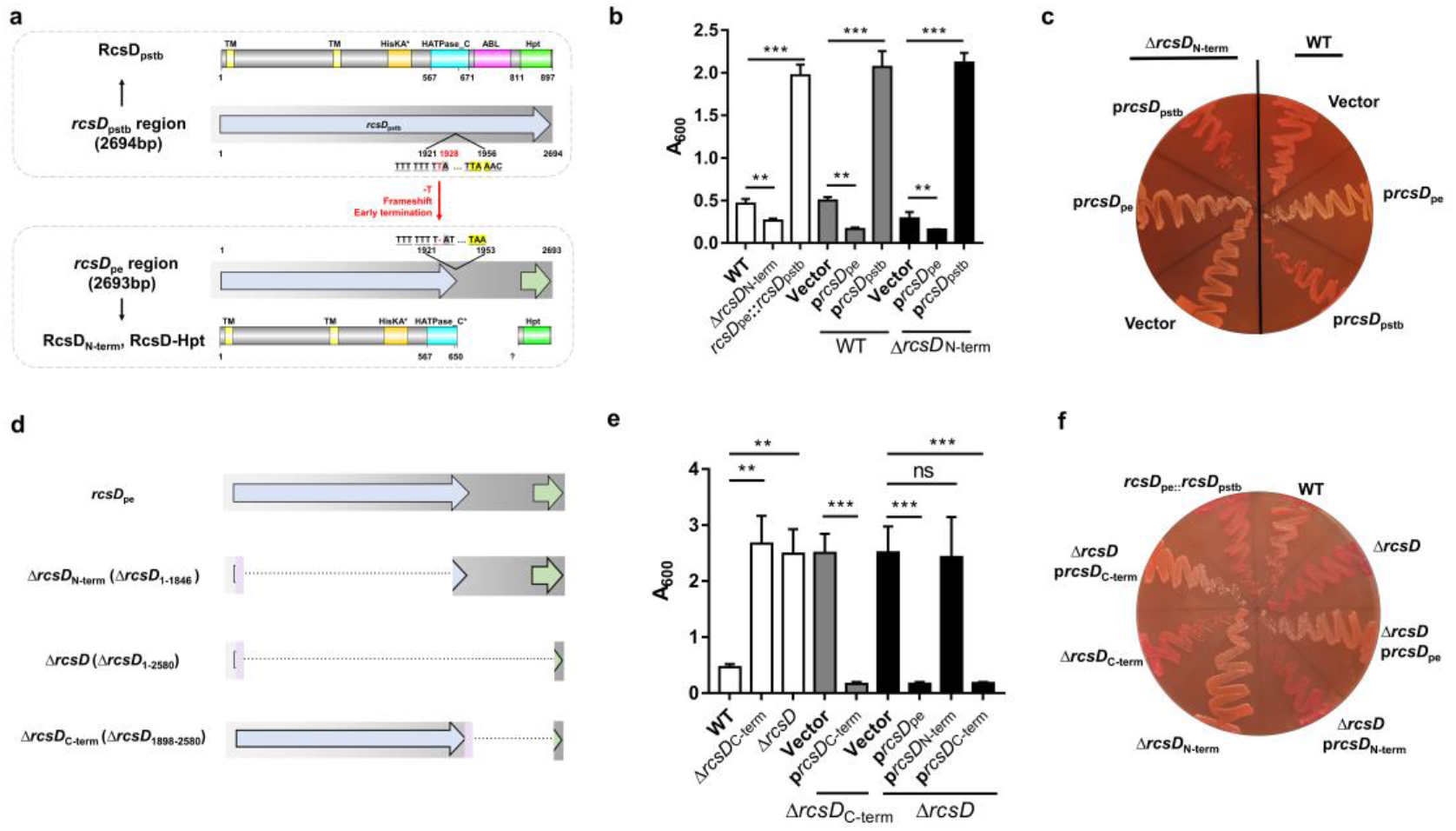
*rcsD*_pe_ negatively regulates biofilm formation, while *rcsD*_pstb_ positively regulates biofilm formation in *Y. pestis*. **a** Schematic representation of the *rcsD* frameshift mutation that occurred during speciation of *Y. pestis* from its ancestor *Y. pseudotuberculosis. rcsDpstb: rcsD* from *Y. pseudotuberculosis; rcsDpe: rcsD* from *Y. pestis;* RcsDN-term: N-terminal fragment of RcsD; RcsD-Hpt: C-terminal HPt domain of RcsD. Crystal violet (CV) binding assay (**b**) and CR pigmentation assay (**c**) using *Y. pestis* KIM6+ (WT), an N-terminal deletion mutant (Δ*rcsD*_N-term_), and an *rcsD*_pstb_ substitution strain (*rcsD*_pe_::*rcsD*_pstb_) and their derivatives that carry plasmids harbouring *rcsD*_pe_ (p*rcsD*_pe_) or *rcsD*_pstb_ (p*rcsD*_pstb_). **d** Schematic representation of the *rcsD* mutations constructed in this study. CV binding assay (**e**) and CR pigmentation assay (**f**) using *Y. pestis* KIM6+ (WT) and its derivative strains harbouring plasmids expressing different truncations of RcsD. CV assays in panel **b** and **e** were performed together. Error bars represent ± SD from three independent experiments with three replicates. Statistical analysis was performed using one-way analysis of variance (ANOVA) with Dunnett’s multiple comparisons posttestt. ns, not significant; *p < 0.05, **p < 0.01, *** p < 0.001.

Next, we constructed two mutants lacking the entire *rcsD*_pe_ gene, and a C-terminal fragment (*ΔrcsD* and *rcsD*_c-term_) (Figure 1d). In contrast to the *rcsD* N-terminal deletion mutant (Δ*rcsD*_N-term_), these two strains showed comparable phenotypes with increased CR adsorption and *in vitro* biofilm formation (Figure 1e and 1f). The opposing effects on biofilm formation in these mutants suggest that the C-terminal region may contribute to the inhibition of biofilm formation. To test this hypothesis, we expressed *rcsD*_c-term_, *rcsD*_N-term_ and *rcsD*_pe_ in an *rcsD* deletion mutant. Overexpression of *rcsD*c-term and *rcsD*_pe_ but not *rcsD*_N-term_ greatly decreased CR pigmentation and biofilm formation (Figure 1e and 1f). Collectively, these results demonstrate that RcsDpstb positively regulates, while the frameshifted *rcsD*_pe_ negatively regulates, biofilm formation, and that this activity depends on the C-terminus.

### *rcsD*_pe_ expresses intact RcsD and a small HPt-containing domain protein RcsD- Hpt

The C-terminal HPt domain of *rcsD* is phosphorylated by RcsC and subsequently transfers a phosphate group to the response regulator RcsB *in vitro* (Takeda et al., 2001). Given that frameshifted *rcsD*_pe_ is functional, and expression of the C-terminus of *rcsD* is sufficient to repress biofilm formation, we hypothesised that a small protein containing HPt domain is expressed by *rcsD*_pe_. To identify the putative open reading frame (ORF) of this latter protein, we analysed candidate translational initiation sites in the C-terminus of *rcsD*_pe_ by searching for AUG, GUG and UUG codons, which account for approximately 80%, 12% and 8% of start codons in bacterial genomes (Srivastava et al., 2016). We found two putative ORFs (encoding genes of 462 and 573 nucleotides, respectively, Supplementary Figure 1a) with UUG start codons, denoted UUG^-462^ and UUG^-573^. Ectopic expression of these two putative ORFs in *Y. pestis* KIM6+ showed comparable biofilm formation to expression of full length *rcsD*_pe_ (Supplementary Figure 1b). Mutation of these two start codons alone (UUG^-462^**→**UUA, UUG^-573^**→**CUU) or together did not alter the function of *rcsD*_pe_ (Supplementary Figure 1c), indicating they are not the start sites promoting expression of the functional protein.

AUU, CUG and AUC are occasionally used as start codons in bacteria (Cao and Slavoff, 2020; D’Lima et al., 2017). We identified a putative ORF, encoding a 103- residue protein, initiated by an AUU codon, which also had a predicted RBS binding site (Figure 2a). Mutation of this codon or the predicted RBS region in *rcsD*_pe_ abolished its function, while replacement of AUU with a strong AUG start codon significantly enhanced its function (Figure 2b and 2c). Furthermore, ectopic expression of the 103-residue HPt domain with an AUU start codon, but not with a GGU codon, using a modified pBAD vector in the *Y. pestis* KIM6+ wild-type strain, or *rcsD* deletion mutant, strongly repressed biofilm formation and CR pigmentation (Supplementary Figure 1d), indicating that AUU is a functional start codon initiating translation of RcsD-Hpt.

**Figure 2.**
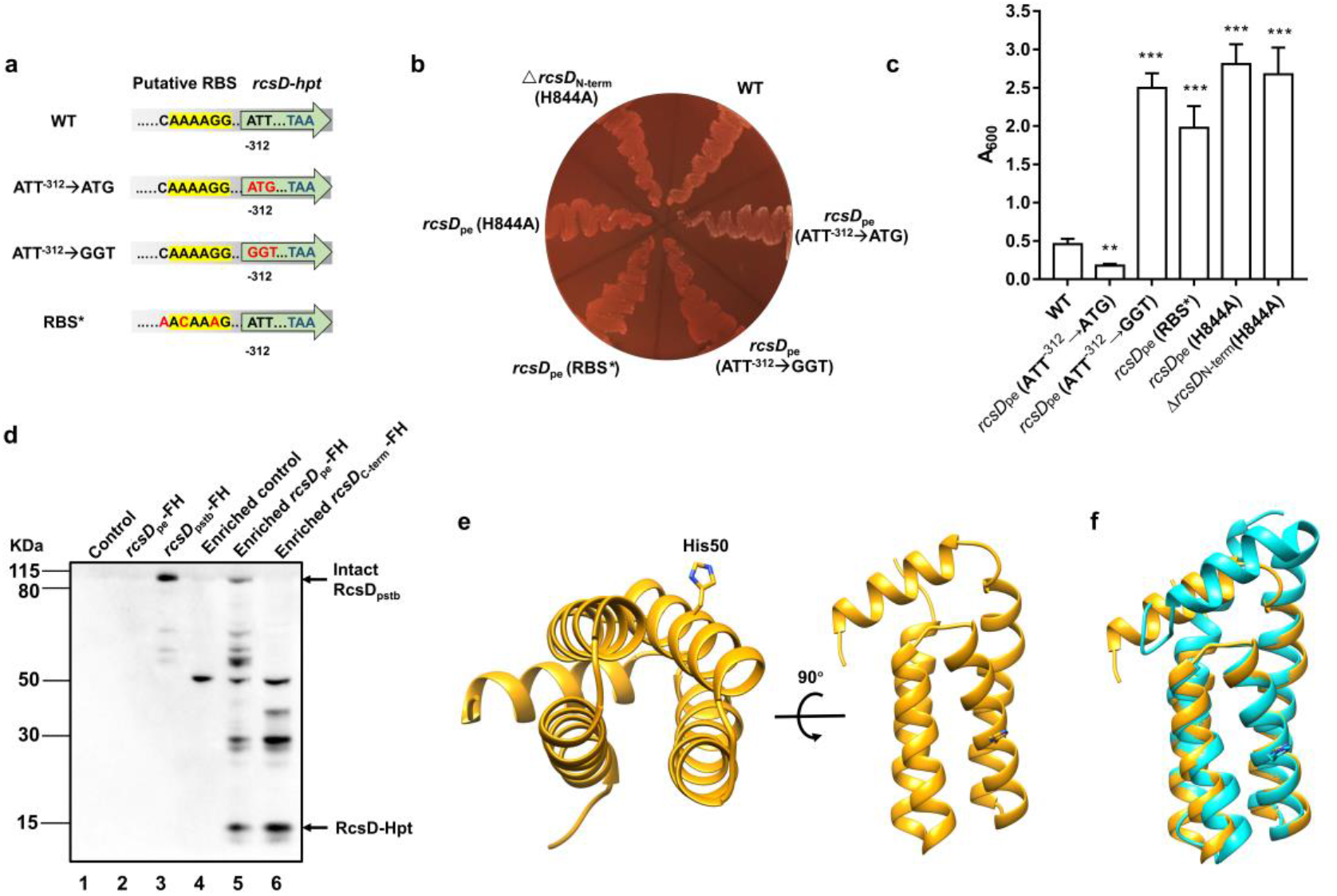
Expression of RcsD-Hpt is initiated by an uncommon AUU start codon and negatively regulates biofilm formation in *Y pestis*. **a** Schematic representation of mutations introduced to the putative RBS and start codon of RcsD- Hpt. ATT^-312^→GGT or ATT^-312^→ATG indicate that the predicted ATT start codon was mutated to GGT or ATG, respectively. RBS* indicates that the putative RBS of RcsD-Hpt was mutated. CR pigmentation assay (**b**) and CV binding assay (**c**) using *Y. pestis* KIM6+ (WT) and its derivative strains. Error bars represent ± SD from three independent experiments with three replicates. Statistical analysis was performed using one-way ANOVA with Dunnett’s multiple comparisons posttest, comparing each construct with WT. *p < 0.05, **p < 0.01, *** p < 0.001. **d** Expression of RcsD in *Y. pestis* and its derivatives were detected by western blot analysis using an anti-Flag antibody. The 3xFlag and His6 epitope tags (FH) encoding sequences were fused to the C-terminus of *rcsD*_pe_ and *rcsDpstb*. **e** Structure of RcsD-Hpt (103 residues) predicted by AlphaFold2. Conserved His50 is located at the a3 helix of RcsD-Hpt. **f** Structure comparison of RcsD-Hpt (103 residues) by AlphaFold2 (yellow) and HptB of *Pseudomonas aeruginosa* PAO1 (PDB 7C1l, cyan).

Next, we introduced 3× Flag and 6× His tags to *rcsD*_pe_ and *rcsD*_pstb_ and analysed their expression by western blotting. A band in the tagged RcsD_pstb_ strain was observed at the expected size (Fig 2d, Lane3), but no visible band was detected in the tagged *rcsD*_pe_ strain (Fig 2d, Lane2). To enhance detection sensitivity, the *rcsD*_pe_ lysates were enriched using Nickel-agarose before western blotting. An approximately 15kDa band, corresponding to the predicted protein containing the HPt domain, was detected in both *rcsD*_pe_ and *rcsD*C-term expressing strains (Fig 2d, Lane 5 and Lane 6). In addition, full length RcsD was detected in *rcsD*_pe_ but not in *rcsD*_c-term_ expressing strains, indicating that intact RcsD is weakly expressed in wild-type *Y. pestis* through translational recoding (Rodnina et al., 2020). This is in accordance with the translational *lacZ* reporter system, which detected 1% readthrough once frameshifted (Supplementary Figure 1e). Taken together, these data suggest that a 103 amino acid protein, designated as RcsD-Hpt, is expressed by *rcsD*_pe_ and functions as a negative regulator of biofilm formation in *Y. pestis*.

RcsD is a phosphorelay protein that can transfer phosphate from the conserved His residue in its HPt domain to a conserved Asp in the receiver domain of RcsB and can also dephosphorylate this site (Ancona et al., 2015; Takeda et al., 2001). An H844A mutation in the *Y. pestis* wild type or *rcsD*_N-term_ deletion strain showed a comparable phenotype to a full *rcsD* deletion strain (Figure 2b, c and Figure 1e), while plasmid-borne expression of the mutated version of *rcsD*_pe_ or *rcsD-hpt* in *Y. pestis* wild type and its *rcsD* mutants displayed a similar phenotype to an empty vector control (Supplementary Figure 1f). Next, we modelled the structure of RcsD-Hpt with AlphaFold2 (Fowler and Williamson, 2022) (Figure 2e), which revealed a similar structure to HptB, an HPt orphan protein in *Pseudomonas aeruginosa (*Figure 2f) (Chen et al., 2020). Like HptB, RcsD-Hpt forms an elongated bundle of four helices α2, α3, α4 and α5, covered by the short N-terminal α1 helix. The imidazole side chain of the conserved active-site histidine residue His50 (His844 in RcsD_pstb_) is located near the middle of helix α3 and protrudes from the bundle where it is exposed, as His57 is in HptB. Taken together, RcsD-Hpt may function as a classical HPt orphan protein, and the conserved His residue is crucial for its function.

### A frameshift in *rcsD* alters Rcs signalling in *Y pestis*

RcsF and IgaA, which regulate environmental stress sensing (Cho et al., 2014; Guo and Sun, 2017; Wall et al., 2018), transfer signals to RcsD through interaction of IgaA with its periplasmic domain (Wall et al., 2020). Given that RcsD-Hpt does not encode a functional periplasmic domain, we hypothesised that the roles of RcsF and IgaA are dispensable in the *Y. pestis* Rcs signalling system. We constructed *rcsF* and *igaA* deletion mutants in the *Y. pestis* wild type and *rcsD*_pstb_ substitution strains, respectively. In the *Y. pestis RcsD_pstb_* substitution strain, deletion of *rcsF* increased CR adsorption and biofilm formation (Fig.3a and 3b), while deletion of *igaA* completely abolished biofilm formation and CR binding (Figure 3c and 3d). Furthermore, overexpression of RcsF but not its C125S mutant, which was reported to be inactive due to the cysteine to serine mutation (Rogov et al., 2011), decreased biofilm formation and CR adsorption (Figure 3a and 3b), while expression of IgaA but not its C413S mutant complemented the phenotype of the *igaA* deletion strain (Pucciarelli et al., 2017) (Figure 3c and 3d). These results indicate that when receiving signals transduced from RcsF and IgaA, RcsD_pstb_ dephosphorylates RcsB, and thus Rcs signalling is switched off. Instead, deletion or ectopic expression of *rcsF* or *igaA* does not regulate biofilm formation and CR pigmentation in wild type *Y. pestis* KIM6+ (Figure 3a-3d), suggesting decoupling of the requirement for RcsF and IgaA and the Rcs system after the transition from *rcsD*_pstb_ to *rcsD*_pe_.

**Figure 3.**
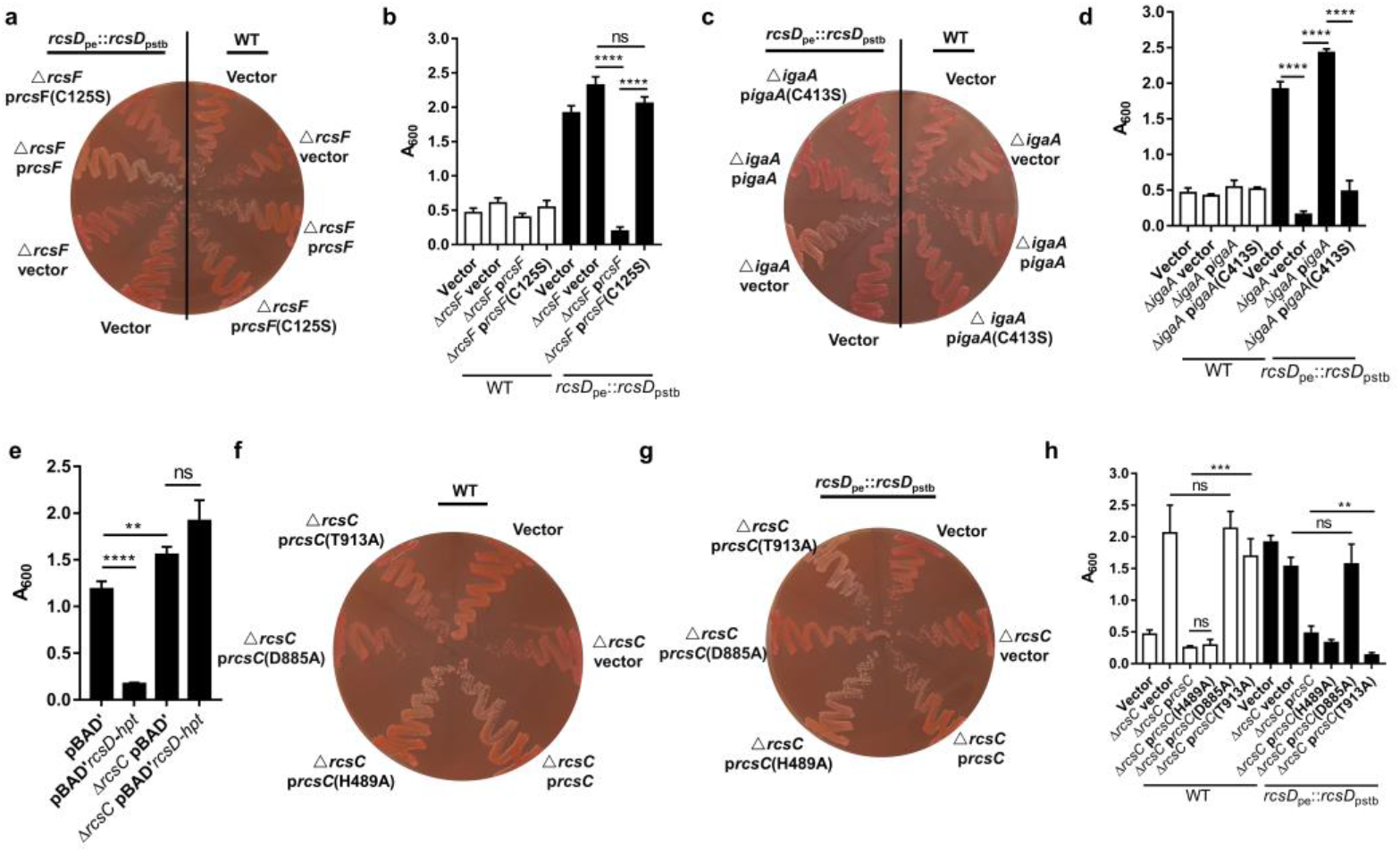
The frameshift mutation in *rcsD* alters the Rcs signalling pathway in *Y. pestis*. CR pigmentation assay (**a** and **c**) and CV biofilm assay (**b** and **d**) the derivatives of *Y. pestis* KIM6+ (WT) and the *rcsD*_pstb_ substitution strain. *prcsF*, plasmid expressing *rcsF; prcsF* (C125S), plasmid expressing *rcsF* with Cysteine (C) to Serine (S) substitution at position 125, *pigaA*, plasmid expressing *igaA; pigaA* (C413S), plasmid expressing *igaA* with Cysteine (C) to Serine (S) substitution at position 413. **e** CV binding assay using WT and an *rcsC* deletion mutant expressing RcsD-Hpt (*rcsD-hpt*). CR pigmentation assay (**f** ang **g**) and CV biofilm assay (**h**) using two *Y. pestis rcsC* deletion mutants expressing different *rcsC* variants. CV assays in panel **b**, **d** and **h** were performed together. Error bars represent ± SD from three independent experiments with three replicates. Statistical analysis was performed using unpaired t test. ns, not significant; *p < 0.05, **p < 0.01, *** p < 0.001, **** p < 0.0001.

RcsC, a bifunctional histidine kinase and phosphatase, phosphorylates and dephosphorylates RcsD, which subsequently, via RcsB, activates or represses expression of its target genes (Guo and Sun, 2017; Wall et al., 2018). Deletion of RcsC resulted in increased biofilm formation and CR pigmentation (Sun et al., 2008) (Figure 3e), indicating RcsC is involved in Rcs signalling in *Y. pestis*. In addition, high expression of RcsD-Hpt in the *rcsC* deletion mutant did not affect biofilm formation and pigmentation (Figure 3e), indicating that RcsC is required for RcsD-Hpt-mediated biofilm repression in *Y. pestis*. His489 and Asp885 in RcsC are the predicted autophosphorylation and subsequent transfer receipt sites, respectively (Clarke et al., 2002; Latasa et al., 2012). A D885A mutation in RcsC abolished its function in an *rcsD*_pstb_ substitute or wild-type strain (Figure 3f-3h), indicating that Asp885 was crucial for phosphate transfer in both conditions. An H489A mutation in RcsC did not alter its function in the wild-type strain (Figure 3f and 3h), indicating that His489 is not important for the phosphorylation of RcsD-Hpt. Surprisingly, mutation of His489 resulted in decreased biofilm formation and CR adsorption in the *rcsD*_pstb_ substitution strain (Figure 3g and 3h), indicating that His489 might be function as a phosphate reservoir involved in dephosphorylation of RcsD. RcsC T903A constitutively activates Rcs in *Salmonella enterica* (García-Calderón et al., 2005; Latasa et al., 2012). The corresponding T913A mutation in RcsC strongly decreased biofilm formation and CR pigmentation in the RcsD_pstb_ substitution strain (Figure 3f-3h), indicating that an RcsC T913A mutant stimulates Rcs in a similar manner to *S. enterica*. The same mutation in RcsC resulted in enhanced biofilm formation in *Y. pestis*, indicating that an RcsC T913A mutant has an impaired ability to phosphorylate RcsD-Hpt. Taken together, these data indicate that RcsC remains a crucial component of the Rcs phosphorelay system in *Y. pestis*.

RcsB is a phosphoacceptor in the Rcs system and contains a conserved Asp site in its receiver domain, which can be phosphorylated by RcsD (Fredericks et al., 2006; Takeda et al., 2001). Phosphorylated RcsB directly regulates expression of biofilm related genes (*hmsT, hmsD, hmsP* and *hmsHFRS*) in *Y. pestis* (Fang et al., 2015; Guo et al., 2015; Sun et al., 2012). Expression of *rcsD*_pe_ or *rcsD*_pstb_ only conferred altered biofilm formation in the presence of RcsB (Figure 4a), indicating that RcsDpe and RcsD_pstb_ might differentially modulate the phosphorylation of RcsB. To test this hypothesis, we detected the phosphorylation of RcsB using Phos-tag^TM^ SDS-PAGE gels and western blotting (Madec et al., 2014). As expected, replacement of *rcsD*_pe_ by *rcsD*_pstb_ in *Y. pestis* KIM6+ resulted in decreased phosphorylation of RcsB, while mutation of the conserved Asp residue (D56Q) in RcsB abolished the phosphorylated protein band (Figure 4b). Consistent with the phosphorylation status of RcsB, the transcription and expression of HmsT were differentially regulated by RcsDpe and RcsD_pstb_ (Figure 4c and 4d).

**Figure 4.**
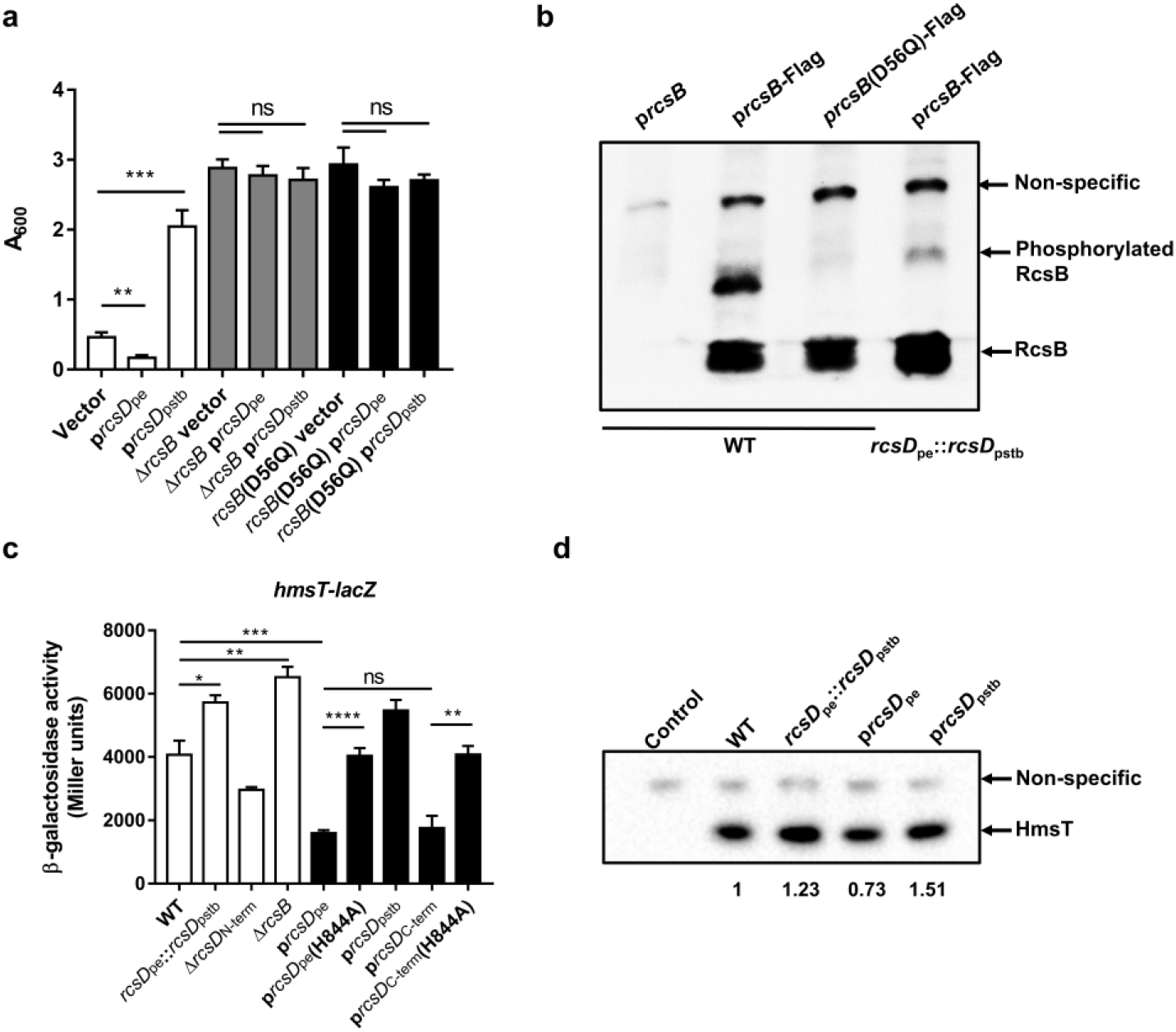
The frameshift mutation in *rcsD* increases the phosphorylation of RcsB and represses the expression of *hmsT*. **a** CV biofilm assays using a WT (white), *ΔrcsB* (grey) or *rcsB* (D56Q) (black)mutant strain, each harboring pUC19 vectors expressing *rcsD*_pe_ or *RcsD_pstb_*. **b** Phosphorylation analysis of RcsB in the *Y. pestis* KIM6+ strain (WT) harbouring an RcsB expression plasmid *(prcsB)*, a plasmid expressing RcsB fused with a 3xflag tag (p*rcsB-*Flag), or a modified p*rcsB*-Flag expression plasmid in which the conserved phosphorylation site Asp50 was mutated to Gln, and the *Y. pestis RcsD_pstb_* substitution strain harbouring the p*rcsB-*Flag expression plasmid. **c** Quantification of *hmsT* expression using a β-galactosidase assay. The LacZ reporter gene was fused with the *hmsT* promoter in plasmid pGD926. **d** Expression of HmsT was analysed by western blotting using an anti-Flag antibody. Error bars represent ± SD from three independent experiments with three replicates. Statistical analysis was performed using one-way ANOVA with Dunnett’s multiple comparisons post-test. ns, not significant; *p < 0.05, **p < 0.01, *** p < 0.001.

### The frameshift mutation in *rcsD* promotes retention of the *pgm* locus during *Y. pestis* flea infection

Loss of function in *rcsA* is a crucial step for *Y. pestis* to establish flea-borne transmission (Sun et al., 2008; Sun et al., 2014). We therefore speculated that mutation of *rcsD* might also play a role in the adaptation of *Y. pestis* to the flea. We therefore infected the Oriential rat flea, *Xenopsylla cheopis*, with *Y. pestis* wild type (KIM6+), the *rcsD*_pe_::*rcsD*_pstb_ wherein *rcsD*_pe_ was substituted with *rcsD*_pstb_ and the *rcsD*_N-term_ deletion strain. Bacterial burdens in infected fleas at 0, 7 and 28 day (s) post infection were not significantly different between any strain combinations (Figure 5a). In addition, we did not observe significant differences in flea blockage, despite different *in vitro* capacities to form biofilms (Figure 5b). Taken together, these results suggest that the frameshift mutation in *rcsD* does not alter the infection, persistence, and blockage-forming capacity of *Y. pestis* in fleas.

**Figure 5.**
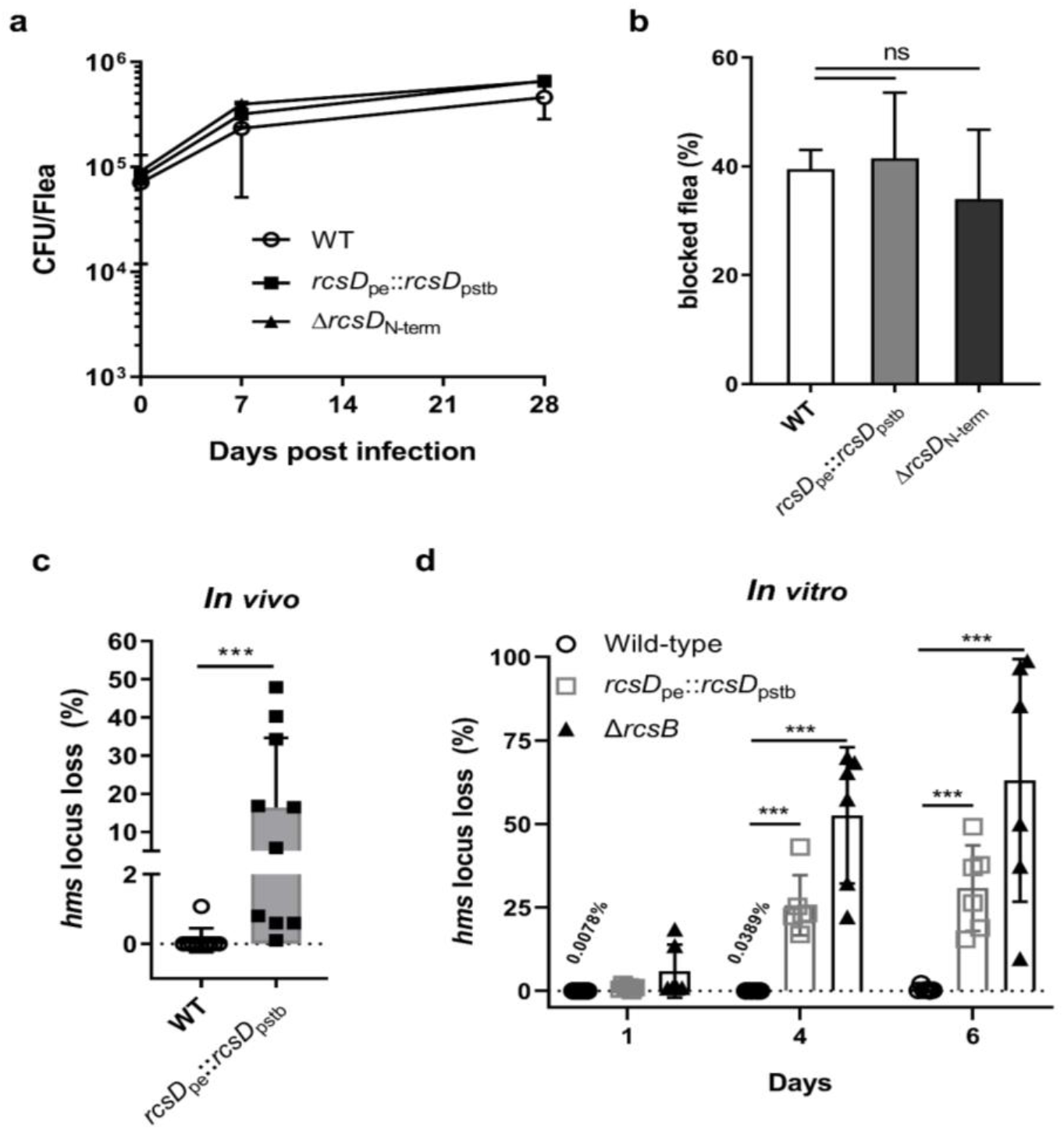
The frameshift mutation in *rcsD* stabilizes the *pgm* locus in *Y. pestis*. **a** Bacterial burdens in fleas infected with *Y. pestis* WT, *rcsD*_pe_::*rcsD*_pstb_ and *rcsD*_N-term_ strains after 0, 7 and 14 days of infection. **b** Cumulative blockage of fleas after 4 weeks of infection with *Y. pestis* WT, *rcsDpe*::*RcsD_pstb_* and Δ*rcsD*_N-term_ strains. Two independent infection experiments are shown. **c** Percent of *pgm* locus loss in fleas infected with *Y. pestis* WT and *rcsD*_pe_::*rcsD*_pstb_ after 4 weeks of infection. Ten infected fleas were used for this assay. Statistical analysis was performed using a Fisher’s exact test. **d** Percent of *pgm* locus loss *in vitro* with *Y. pestis* WT, the *rcsD*_pe_::*rcsD*_pstb_ substitution strain and the *ΔrcsB* strain. One-way ANOVA with a Dunnett’s post-test were performed for statistical analysis. Error bars represent ± SD from three independent experiments with six replicates. ns, not significant; *p < 0.05, **p < 0.01, *** p < 0.001.

We fortuitously observed that a *Y. pestis rcsD*pstb strain, but not the wild type, displayed a *pgm*- phenotype after 4 weeks of flea infection. Prior to the infectious blood meal, we observed no *pgm*- phenotype when the wild-type and *rcsD*_pstb_ samples were plated on CR agar plates. Twenty-eight days post infection, we observed the *pgm-* phenotype in bacteria isolated from all fleas infected with the *rcsD*_pe_:: *rcsD*_pstb_ swapped strain. The mean percentage of isolates displaying the *pgm-* phenotype was 16.3%, ranging from 0.1% to 47.9%, whereas only one out of ten fleas infected with wild-type strain displayed this phenotype, with a frequency of ~1.0% (Figure 5c). We further analysed the *pgm* mutation rate of *Y. pestis* in liquid medium. To mimic the environment in the flea, *Y. pestis* were grown in liquid medium for several days, where they remained in stationary phase before reinoculation. Consistent with the flea infection data, *Y. pestis* encoding *rcsD*_pstb_ displayed a significantly higher *pgm* mutation rate relative to the wild-type strain (Figure 5d). PCR analysis confirmed that the *pgm-* phenotype was caused by the spontaneous deletion of 102 kb of *pgm* locus (Fetherston et al., 1992) (Tong et al., 2005) (data not shown). Taken together, these results suggest that the frameshift mutation in *rcsD* promoted stable maintenance of the *pgm* locus of *Y. pestis* KIM6+ in infected fleas.

### Genome-wide identification of genes regulated by the Rcs phosphorelay system in *Y. pestis*

Rcs has been reported to modulate virulence in other bacteria (Li et al., 2015; Wall et al., 2018; Wang et al., 2012). KIM6+ is an avirulent derivative of the fully virulent KIM strain, which was cured of the pCD1 plasmid. To investigate the role of Rcs on pathogen virulence in a mammalian host, we took advantage of the *Y. pestis* biovar Microtus strain 201 (Zhang et al., 2014), a human-avirulent but rodent-virulent strain, isolated from a natural reservoir, the Brandt’s vole *(Microtus brandti)* (Song et al., 2004). Mutation of *rcsB* or *rcsD* in *Y. pestis* biovar Microtus strain 201 showed similar CR absorption and *in vitro* biofilm phenotype as *Y. pestis* KIM6+ strain (data not shown). To our surprise, deletion of *rcsB* or replacement of *rcsD*_pe_ by *rcsD*_pstb_ in this strain did not significantly affect their virulence when mice were infected with different doses of bacteria subcutaneously (Figure 6a-c).

**Figure 6.**
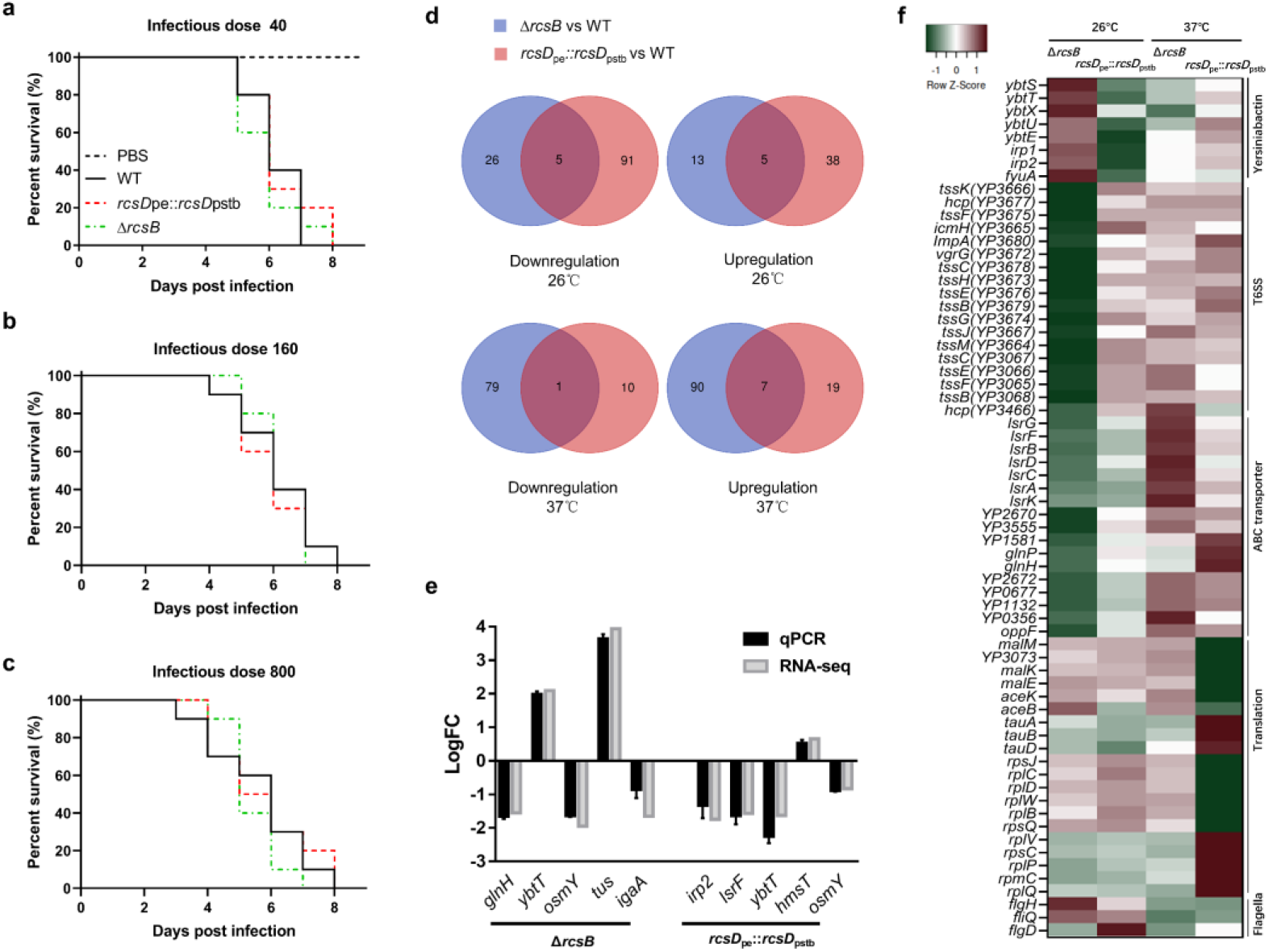
Genome-wide identification of genes regulated by the Rcs phosphorelay system in *Y pestis*. **a-c** Survival of C57BL/6 mice infected with *Y. pestis* Microtus strain 201 and its derivatives using an infectious dose of 40, 160 and 800 CFU. **d** Venn diagram of upregulated and down regulated genes in *Y. pestis* strains growing at temperatures (26°C or 37°C). **e** qPCR analysis of *hmsT* and differentially expressed genes (DEGs) identified by RNA-seq. The screening threshold for DEGs was defined as ļlogFCļ ≥ 1, and p ≤ 0.05). Error bars indicate SD from at least three samples. **f** Heatmaps showing the differential expression of genes identified through COG analysis.

Rcs has been reported to modulate the expression of many genes in response to environmental stress. To characterize the genes regulated by the Rcs system in *Y. pestis*, we performed RNA-seq on total RNA isolated from *Y. pestis* biovar Microtus strain 201, *rcsB* deletion and *rcsD* substitution strains cultured at 26°C and 37°C. A total of 139 genes (43 upregulated and 96 downregulated) and 49 genes (18 upregulated and 31 downregulated) were significantly differentially expressed (ļlogFCļ ≥ 1, p ≤ 0.05) in the RcsB mutant and RcsD substitute strain when compared with the wild-type strain at 26°C (Figure 6d, Supplementary Figure 2a-b, and Supplementary Table 4). At mammalian temperature (37°C), 37 genes (26 upregulated and 11 downregulated) and 177 genes (97 upregulated and 80 downregulated) were significantly differentially expressed (Figure 6d, Supplementary Figure 2c-d and Supplementary Table 4). Several differentially expressed genes identified by RNA-seq were verified by qRT-PCR, indicating comparable patterns of expression (Figure 6e, and Supplementary Table 4). Substitution of *rcsD*_pstb_ had an opposing effect on gene expression to deletion of *rcsB* (Figure 6f, and Supplementary Table 4), indicating that the frameshift mutation in *rcsD* lessens the regulatory function of Rcs. Furthermore, Rcs positively regulated genes such as those encoding the Type 6 Secretion System (T6SS), those related to biosynthesis of yersiniabactin and ABC transporter genes (Figure 6f). These observations suggest that Rcs might play an important role for the environmental fitness and virulence of *Y. pestis*.

### A frameshift mutation in *rcsD* is an evolutionary step present in modern *Y. pestis* lineages

To investigate the role of *rcsD* mutation in the evolution of *Y. pestis*, we analysed the evolutionary changes that occurred during the divergence of *Y. pestis* from *Y. pseudotuberculosis (*Figure 7 and Supplementary Table 5). Mutation of *rcsD* is present in all *Y. pestis* that harbor five genetic changes *(pde3’, ymt, rcsA, pde2* and *ureD)* required for flea colonization (Figure 7), except for the 0.PE7 branch (Figure 7). An IS element in the *ompC* gene is one of two IS elements driving instability of the *pgm* locus (Figure 7 and Supplementary Table 5) (Fetherston et al., 1992; Tong et al., 2005). The IS element in *ompC* was present in most modern *Y. pestis* branches, but not in RT5, an ancient *Y. pestis* strain isolated from a bubonic plague patient during the Bronze Age (Spyrou et al., 2018). This indicates that emergence of the *rcsD* mutation is likely not due to loss of the *pgm* locus caused by IS elements. Although multiple *rcsD* mutations are present across the phylogeny (Figure 7 and Supplementary Table 5), the HPt encoding region is present in all sequenced *Y. pestis* isolates, indicating an important role of RcsD-Hpt in refining stable blockage-mediated flea-borne transmission of *Y. pestis*.

**Figure 7.**
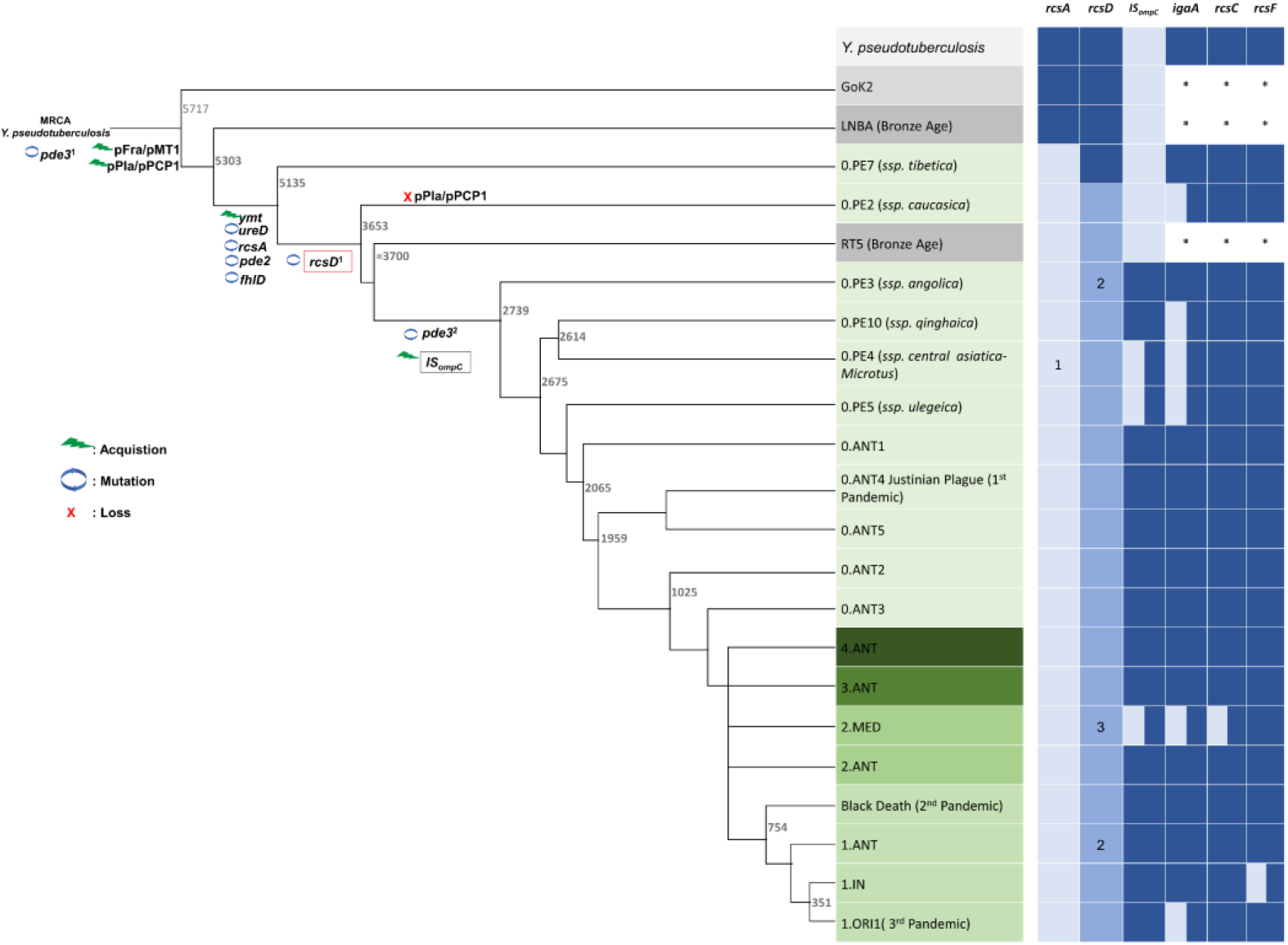
Genetic changes in Rcs genes during speciation of *Y. pestis*. Acquisition events are indicated by green lightning symbols, loss of genetic material by a red cross and mutation by blue circles. All comparisons shown are relative to *Y. pseudotuberculosis*. Increasingly darker shades of grey represent *Y. pseudotuberculosis*, and ancient strains from the Iron Age and the LNBA lineage respectively. Increasingly darker shades of green represent branches one to four as annotated in the figure. Rcs-related genes and IS*ompC* were considered as intact (dark blue), mutated (light blue), or absent (white). *: not analyzed. 1, present in all *Y. pestis* except branch 0 strain 0.PE4b; 2, *rcsD*_N-term_ and *rcsD*C-term were located in a different genome site in the Nairobi (1.ANT), Angola (0.PE3) and Algeria3 (ORI) strains due to chromosome rearrangement; 3, indels are present in *rcsD*_N-term_ in some strains I-3086 (0.PE4m) and A-1825 (2.MED1). MRCA, The most recent common ancestor. The figure and nomenclature are adapted from Demeure, et al., 2019 (Demeure et al., 2019).

## Discussion

*Y. pestis* and its ancestor *Y. pseudotuberculosis* have historically been studied as models for pathogen evolution, and have helped to shape our understanding of the evolutionary processes driving niche adaptation, transmission and pathogenesis (Wren, 2003). A recent paleogenomic study has clarified major steps driving evolution of *Y. pestis* (Rasmussen et al., 2015; Spyrou et al., 2018). An ancestral *Y. pseudotuberculosis*, which has a mutation in the promoter region of *pde3* (Sun et al., 2014), acquired two plasmids, pPla and pMT1, in addition to other genetic changes, becoming the ancient virulent *Y. pestis* (Bearden et al., 2009; Cui et al., 2013; Sodeinde et al., 1992; Zimbler et al., 2015). At this point, *Y. pestis* may still have been prevalent in the environment, where it could be transmitted to humans and animals by the faecal-oral route and occasionally by flea bites through early phase transmission (Supplementary Figure 3). Later, other genetic changes, including mutations in *ymt*, *rcsA*, *pde2* and *ureD*, occurred in the ancient *Y. pestis*, fully converting the pathogen to a flea transmitted modality (Chain et al., 2004; Cui et al., 2013; Hinnebusch, 2005) (Supplementary Figure 3).

Compared to its ancestor, which faced multiple changing environments, the establishment of a flea-mammalian host transmission cycle limited environmental exposure of *Y. pestis*. The progenitor lineage of flea-borne *Y. pestis* still required a series of genetic changes to repurpose its environmental signal sensing and transduction systems to adapt to its new lifestyle and niche (Laayouni et al., 2014). Loss of redundant genes and response pathways may have contributed to the fitness of *Y. pestis*. For example, mutation of flagella-related genes in *Y. pestis* occurred in parallel to its colonisation of the flea (Minnich and Rohde, 2007). Analysis of a broad set of *Yersinia* genomes demonstrated that deletion of one thymine in *rcsD* occurred after the ancient *Y. pestis* acquired the major genetic changes required for flea-borne transmission (Figure 7 and Supplementary Figure 3). The frameshift mutation in *rcsD* might compensate for the fitness cost imposed by loss of *rcsA* function by dampening the subsequent drastic changes in gene expression.

The frameshift present in *rcsD*_pe_ leads to the expression of two functional proteins: RcsD-Hpt and intact RcsD. Low levels of intact RcsD may be expressed by translational readthrough, while RcsD-Hpt is expressed from a rare AUU start codon. Intact RcsD and RcsD-Hpt have different functions in *Y. pestis*. RcsD dephosphorylates RcsB, while RcsD-Hpt phosphorylates RcsB. This subsequently promotes different capacities for biofilm formation and likely multiple other phenotypes. The periplasmic domain of RcsD receives environmental signals sensed by RcsF and IgaA, which in turn regulates the phosphorylation of RcsD by RcsC (Wall et al., 2018; Wall et al., 2020). RcsD-Hpt lost the ability to respond to environmental signal transduction by RcsF and RcsD but could still receive a phosphate group from RcsC. Although RcsD-Hpt and intact RcsD were expressed in *Y. pestis*, RcsD-Hpt appears to play a dominant role in regulation of the Rcs pathway. This hypothesis is supported by two observations: 1) expression of *rcsD*_pe_ conferred a similar phenotype as expression of *rcsD-hpt*, and 2) RcsF and IgaA modulate Rcs signalling in the *rcsD*_pstb_ substitution strain but not in wild-type *Y. pestis*.

Although *rcsD*_pstb_ has the same RBS and start codon, only intact RcsD was detected by western analysis, indicating intact RcsD plays a major role in the background of *rcsD*_pstb_. Sequence analysis indicated that a putative start codon and RBS are present in *rcsD* in many organisms (Supplementary Table 6). This indicates that RcsD-Hpt may play a moonlighting role in Rcs signalling. Indeed, wild-type RcsD in *E. coli* produces low levels of a short phosphotransfer protein (Rogov et al., 2004; Wall et al., 2020). HPt orphan proteins function as phosphate transfer components in multiple phosphorelay systems in numerous prokaryotes and eukaryotes (Herivaux et al., 2018; Kennedy et al., 2016; Mohanan et al., 2017; Valentini et al., 2016), and have evolved from larger phosphotransferase proteins containing multiple domains. The frameshift present in *rcsD* of *Y. pestis* may represent an ongoing evolutionary process generating an orphan HPt protein and consequently a new regulatory pathway.

The *rcsD* frameshift alters the Rcs signalling pathway, which in turn decreases *Y. pestis* biofilm formation. The *rcsD* frameshift in *Y. pestis* does not significantly affect mammalian virulence and flea colonisation in our study, but it may promote bacterial fitness during multiple flea-mammalian host transmission cycles. Deletion of a 102 kb *pgm* locus occurs spontaneously due to presence of two IS elements around this region and the mutation rate is increased with enhanced biofilm formation (Fetherston et al., 1992; Podladchikova et al., 2002; Silva-Rohwer et al., 2021) (Figure 5d). Likewise, our work shows that increased biofilm formation in an *rcsD* mutant promotes biofilm formation with a subsequent increase in *pgm* loss (Silva-Rohwer et al., 2021). The *pgm* locus contains the *ybt* operon, which is involved in iron acquisition and is required for virulence of *Y. pestis* (Fetherston et al., 2010; Sebbane et al., 2010). Indeed, *Y. pestis* strains lacking the *pgm* locus are avirulent and have been used as live plague vaccines in some countries (Bearden and Perry, 1999; Podladchikova et al., 2002). Its notable that prolonged exposure to biofilm stimulating conditions in the flea gut and during *in vitro* growth resulted in loss of the *pgm* locus here. This may explain why no effect on virulence was noted for the *rcsD*_pe_:: *rcsD*_pstb_ strain in our mouse infection studies which utilized strains cultured at 37°C overnight. Enhanced biofilm production through the *rcsA* mutation in the absence of accompanying *rcsD* mutation in ancient *Y. pestis* strains likely caused a high rate of *pgm* locus loss during flea infection thus conferring vaccine-like protection when transferred to the mammalian host. Eventually this would break the transmission cycle between flea and mammalian host. In modern lineages the *rcsD* mutation therefore serves to dampen biofilm production without obvious compromise to flea blockage and infection rates. Thus, stable maintenance of the *pgm* locus required for amplified infection in the mammalian host that can perpetuate the flea-mammal transmission cycle and intensity of plague outbreaks is promoted (Figure 7). Finally, the frameshift mutation of *rcsD* might represent an important step in the emergence of extant ubiquitous lineages of *Y. pestis*.

## Acknowledgments

We thank Dr. Xiaoyun Pang at Institute of Biophysics, Chinese Academy of Sciences (Beijing, China) for help to predict the structure of RcsD-Hpt by AlphaFold2. This work was supported by NSFC (Project 31700072, 31670139 and 31800120), the Non-profit Central Research Institute Fund of Chinese Academy of Medical Sciences (Project 2019HY310001), the CAMS Innovation Fund for Medical Sciences (CIFMS) (Project 2021-I2M-1-043), and NIH R01AI117016-01A1 to VV.

## Author Contributions

Xiao-Peng Guo, performed majority of the experiments, Conceptualization, Validation, Data Curation, Visualization, Formal Analysis, Investigation, Funding acquisition, Writing – part of original draft and Writing-Review & Editing; Hai-Qin Yan, performed flea experiments, Conceptualization, Validation, Data Curation, Visualization, Formal Analysis, Investigation, Funding acquisition, Writing – most of original draft, and Writing-Review & Editing; Wenhui Yang, performed murine-infection experiments, Conceptualization, Validation, Data Curation, Visualization, Formal Analysis, and Investigation; Zhe Yin, Formal Analysis; Viveka Vadyvaloo, Conceptualization, Resources, Funding acquisition, Supervision, Writing-Review & Editing; Dongsheng Zhou, Conceptualization, Resources, Supervision, Writing-Review & Editing, Project administration; Yi-Cheng Sun, Conceptualization, Resources, Funding acquisition, Supervision, Writing – part of original draft, Writing-Review & Editing, Project administration. All authors reviewed the manuscript.

## Ethics Statement

The animal study of flea blockage, related to Figure 5a, 5b and 5c, was performed in strict accordance with the U.S. National Institutes of Health (NIH) Guide for the Care and Use of Laboratory Animals (National Research Council Committee for the Update of the Guide for the and Use of Laboratory, 2011) and as approved by the Washington State University Institutional Animal Care and Use Committee, under the Animal Subject Approval Form (ASAF) 6641 and 6396.

The animal study of murine infection, related to Figure 6a, 6b and 6c, was performed in strict accordance to the Guidelines for the Welfare and Ethics of Laboratory Animals of China and all the animal experiments were approved by the Institutional Animal Care Committee (IACUC) of Military Medical Sciences (AMMS), ethical approval number IACUC-DWZX-2021-057.

## Data availability

All data is available within the paper, its Supporting Information files, and the NCBI GenBank. RNA-seq sequencing data can be accessed in NCBI GenBank using BioProject ID: PRJNA876755.

## STAR+Methods

### Key resources

#### Bacterial strains and plasmids

This study for utilized *Y. pestis* strain KIM6+, which derives from the sequenced strain KIM strain (Deng et al., 2002), but is cured of the pCD1/pYV plasmid required for mammalian virulence, and is competent for flea blockage (Hinnebusch et al., 1996) and biofilm formation (Darby et al., 2002; Sun et al., 2008). Studies in mice utilized the biovar Microtus strain 201 which wis avirulent to human but fully virulent to mouse (Zhang et al., 2014).

Deletion of *rcsC, rcsF* and *igaA* was achieved using a one-step method to integrate PCR products into the chromosome with pKD46, as previously described (Datsenko and Wanner, 2000). Double mutants were made by the sequential application of pKD46- mediated deletion. CRISPR-Cas12a-Assisted Recombineering were used to introduce point mutations, deletions, insertions, and gene replacements in this study (Yan et al., 2017). All strains were verified by PCR, DNA sequencing and plasmid complementation.

For construction of the *hmsT*::*lacZ* reporter, 350 bp of *hmsT* upstream sequence, together with the first seven codons of the *ORF*, were amplified by PCR using KIM6+ chromosome DNA as the template. The DNA fragments were digested with HindIII and BamHI restriction enzymes and cloned into pGD926 (Ditta et al., 1985; Vieille and Elmerich, 1990), generating in plasmids pYC593 and pYC287. Plasmid expressing *rcsC* (D885A), *rcsC* (T913A), *rcsD* (H844A), *rcsD*::*rcsD-3xflag* and *rcsD- 3xflag*::*rcsD-3xflag-his6* were generated by overlapping PCR as described previously (Li et al., 2015).

For inducible *rcsD* expression, the gene was cloned downstream of the arabinose-inducible promoter of plasmid pBAD/Myc-His (Invitrogen). The plasmid used for determining readthrough was generated by cloning a partial sequence of *rcsD* (159 bp for *rcsD*_pstb_ and 158 bp for *rcsDpe*) containing the 8T (frameshifted region) into pMal- *lacZ*, generating plasmids *pMal-RcsD_pstb_-lacZ* and *pMal-rcsD_pe_-lacZ*. An additional stop codon was introduced into the 158 *rcsD*_pe_ sequence of *pMal-rcsDpe-lacZ*, generating plasmid pMal-*rcsD*_pe_-stop-*lacZ*.

All strains and plasmids used in this study are shown in Supplementary Table 1 and 2. Oligonucleotides used in this study are shown in Supplementary Table 3.

## Methods details

### *In vitro* biofilms

Microtiter plate biofilm assays were performed as previously described (Sun et al., 2012). Briefly, bacteria were cultured overnight in LB broth supplemented with 4 mmol CaCl2 and 4 mmol MgCl_2_. Cultures were subsequently diluted into 96-well plates and incubated with shaking for 24 h at 26°C. The wells were washed, and the adherent biofilm was stained with crystal violet, solubilized with 80% ethanol and 20% acetone and measured by *A*_600_. Results from three independent experiments with three technical replicates per experiment.

### β-galactosidase assays

β-galactosidase activities were measured as previously described (Guo et al., 2015; Sun et al., 2012). Briefly, overnight cultures of *Y. pestis* harboring *lacZ* reporters were diluted to an OD_600_ of 0.05 and grown in LB broth at room temperature to an OD_600_ of 1.5. The active beta-galactosidase, encoded by the LacZ reporter gene in *Y.pestis* strains, could cleave o-nitrophenyl-β-D-galactopyranoside (ONPG) substrate to a bright yellow product. The cells were lysed and ONPG solution were added. After incubation at 37°C, the reaction could be stopped by adding 1M Na_2_CO_3_, then absorbance was measured at 420nm. Results were normalized against cell density and incubation time, and shown in Miller units (H., 1972). At least two independent experiments with technical triplicates were performed.

### Quantitative real time PCR

qRT-PCR was carried out as previously described (Sun et al., 2011). Briefly, cells were first grown in LB broth overnight before diluting to an OD_600_ 0.05 in LB and incubating at room temperature to an OD_600_ of 0.8. Total RNA was isolated using the Rneasy Mini Kit (Qiagen). Residual DNA was removed by treatment with rDNase I (Ambion) and confirmed by PCR. cDNA was synthesized from the RNA and used for quantitative PCR on an Applied Biosystems unit (Quant Studio TM 5). The quantity of mRNA was normalized relative to the reference gene *16sRNA* (YP_r1). The relative mRNA expression levels in each strain were normalized to the wild-type samples. Primers and probe sets used in this study are listed in Supplementary Table 3. Results from three independent experiments performed in technical triplicate were analyzed by one-way ANOVA with Bonferroni’s test.

### Western blotting

Western blotting was performed as previously described (Ren et al., 2017). For detection of enriched RcsDpe, *Y*. *pestis* strains expressing with 3xFlag-His6-tagged recombinant genes were grown at 26°Co to stationary phase. Cells were harvested by centrifugation and disrupted by sonication. The protein was enriched 100-fold by Ni-nitrilotriacetic acid (NTA) His resin for western blot analysis if necessary. For detection of HmsT, *Y*. *pestis* strains harboring a plasmid expressing HmsT with 3xFlag were grown at 26°C to stationary phase. Cells were harvested by centrifugation and disrupted by sonication. Approximately 20 ng proteins were loaded for detection of HmsT. The proteins were separated on 10% SDS-PAGE gels transferred to PVDF membranes (Millipore), analyzed by immunoblotting with an anti-Flag antibody produced by Invitrogen (Catalog number: MA1-91878-HRP; RRID: AB_2537626) and detected with ECL Western Detection Reagents (BioRad). Resulting bands were quantitated by densitometry using NIH ImageJ (Gallo-Oller et al., 2018).

### Phos-tag™ SDS-PAGE

For detection of protein phosphorylation, acrylamide gel was mixed with 25 μM Phostag acrylamide (AAL-107, Wako) and 25 μM MnCl2 (Madec et al., 2014). Cells grown to OD_600_ 0.8 were centrifuged and pellets were resuspended in PBS containing Protease Inhibitor Cocktail (Roche) and then lysed with a sonicator. Samples were quickly loaded onto gels containing 25 μM Phos-tag acrylamide and 25 μM MnCl2 prepared and run. After a 10 min wash with WB transfer buffer supplied with 1 mM EDTA, followed by a 10 min wash with transfer buffer without EDTA. Proteins were transferred to PVDF membrane and blotted using an anti-Flag antibody produced by Invitrogen (Catalog number: MA1-91878-HRP; RRID: AB_2537626).

### RNA-seq

Total RNA was extracted using the RNeasy Kit (Qiagen). RNA sequencing and expression quantification were performed by Genewiz. Gene expression levels were further normalized using the fragments per kilobase of transcript per million (FPKM) mapped reads method to eliminate the influence of different gene lengths and sequencing depth (Wang et al., 2009). The edgeR package was used to identify differentially expressed genes (DEGs) across samples with fold changes ≥ 2 and a false discovery rate-adjusted p (P-value) ≤ 0.05 (Anders and Huber, 2010; Robinson et al., 2010). DEGs were then subjected to an enrichment analysis of GO function and KEGG pathways (Harris et al., 2004; Kanehisa and Goto, 2000).

### Murine infection

Animals were handled in strict accordance to the Guidelines for the Welfare and Ethics of Laboratory Animals of China and all the animal experiments were approved by the Institutional Animal Care Committee of Military Medical Sciences. Bacterial cultures at 37°C were washed twice with PBS (pH 7.2) and then subjected to serial 10-fold dilutions with PBS. Dilutions were plated onto Brain Heart Infusion (BHI) agar plates to calculate the numbers of colony-forming units (CFU). For each strain, different doses of bacterial suspension were inoculated subcutaneously at the inguinal region of ten female BALB/c mice (aged 6–8 weeks old), which were obtained from Charles River Laboratories (Beijing, China). Survival was monitored at regulate intervals, and a survival curve was generated with GraphPad Prism 5.0. P-values were determined using the log-rank (Mantel-Cox) test and the Gehan-Breslow-Wilcoxon test; *p* < 0.01 was considered statistically significant.

### Flea blockage

Flea infections and blockage analysis were carried out as previously described (Silva- Rohwer et al., 2021; Sun et al., 2014). *Y. pestis* strains were grown overnight in 3 mL HIB at 26°C with shaking, then diluted into 100 mL HIB to cultivate at 37°C without shaking. The following day harvested cells were suspended in sterile PBS, and optical density at absorbance of 600nm was determined. A commercial preparation of heparinized mouse blood (BioIVT, New York) was inoculated with *Y. pestis* to a final concentration of CFU/mL of ~ 5×10^8^ to 1×10^9^. *X. cheopis* fleas were allowed to feed on the infected blood through a mouse skin membrane. Studies with mice were performed in strict accordance with the U.S. National Institutes of Health (NIH) Guide for the Care and Use of Laboratory Animals (National Research Council Committee for the Update of the Guide for the and Use of Laboratory, 2011) and as approved by the Washington State University Institutional Animal Care and Use Committee.

Mice: A CD-1 mouse breeding colony originally sourced from Envigo (https://www.envigo.com/model/hsd-icr-cd-1) is maintained at WSU Males and females are used. Neonates between the ages of 2-6 days old are used for feeding fleas for breeding and to maintain infected fleas.

Fleas: Xenopsylla cheopis fleas are maintained at WSU since 2010 in Dr Vadyvaloo’s lab. These fleas were originally sourced from Joseph Hinnebusch’s lab at the NIH. Male and females are used for experiments.

### Analysis of pigmentation

For *in vivo* pigmentation phenotype detection, 10 fleas infected with *Y. pestis* WT or the *rcsD*_pe_::*rcsD*_pstb_ substitution strain were collected at the end of the infection period (T = 28 days). Fleas were individually triturated in sterile PBS, and the fractions were serially diluted and cultivated on a CR plate. Two days later, CFU enumeration relevant to the pigmentation phenotype was performed. For *in vitro* pigmentation assays, fresh *Y. pestis* KIM6+ wild type, the *rcsD*_pstb_ substitution and the Δ*rcsB* strains were inoculated into LB medium and grown at 26°C for 24 h. Cultures were diluted 1:100 into fresh LB medium every 3 days. The cultures at 0, 1, 4 and 6 days postinoculation were plated on CR plates for analysis. The loss of the *pgm* locus was confirmed by PCR using primers described in Supplementary Table 3.

### Quantification and statistical analysis

Figure legends detail the quantification and statistical analyses. We conducted the statistical analyses by Graphpad Prism.

