## Supplemental file for "A frameshift in *Yersinia pestis rcsD* leads to expression of a small HPt variant that alters canonical Rcs signalling to preserve flea-mammal plague transmission cycles"

### Supplementary Figures and Tables

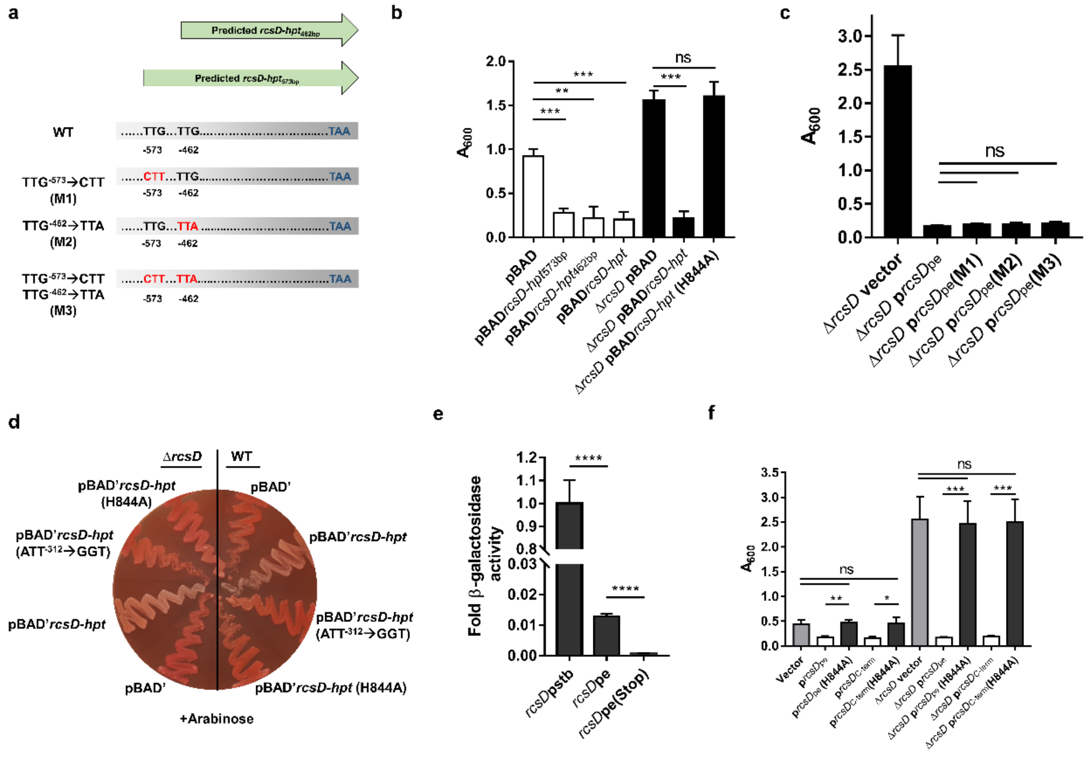


**Supplementary Figure 1. Characterization of proteins encoded by *rcsD*$_{pe}$ region, related to Figure 2.**

**a** Schematic representation of mutations introduced to the putative but non-functional start codons. **b** CV biofilm assay using WT and Δ*rcsD* strains complemented by the expression of putative *rcsD-hpt* genes. **c** CV biofilm using Δ*rcsD* strains complemented by the expression of *rcsD-hpt* with mutated start codons or the mutated conserved phosphorylation site. **d** CR pigmentation assay of ectopic expression of the 103-residue HPt domain with an AUU start codon or GGU codon, using a modified pBAD vector (without ATG codon) in the *Y. pestis* KIM6+ wild-type strain, or *rcsD* deletion mutant. **e** Quantification of readthrough efficiency using a LacZ reporter assay. **f** CV assay of *Y. pestis* expressed with *rcsD* and its H844A-relevant derivatives. CV assays in panel **c** and **f** were performed together. Error bars represent ± SD from three independent experiments with three replicates. Statistical analysis was performed using one-way ANOVA with Dunnett's multiple comparisons posttest. ns, not significant; *p < 0.05, **p < 0.01, *** p < 0.001.

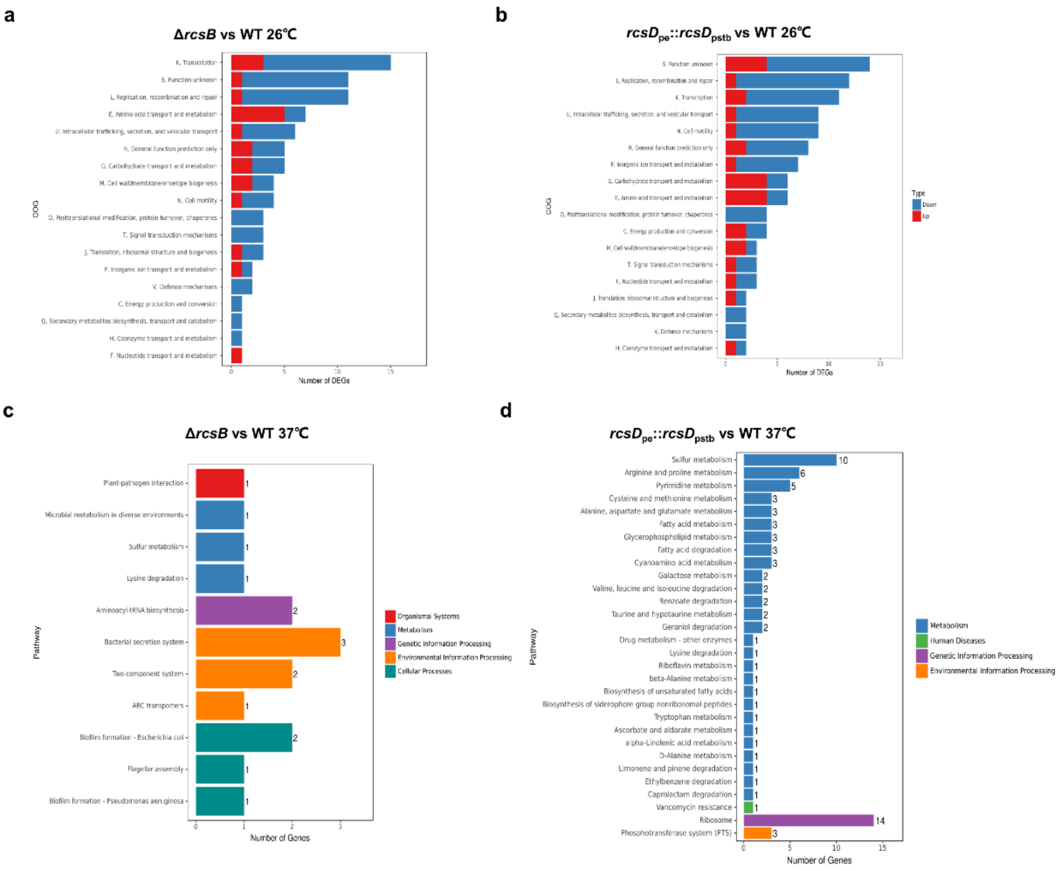


**Supplementary Figure 2. RNA-seq analysis of genes regulated by Rcs system, related to Figure 6.**

**a-d** Clusters of orthologous groups (COG) analysis of RNA-seq gene expression data comparing the Δ*rcsB* strain and WT (**a**), and $rcsD_{pe}$::$rcsD_{pstb}$ with WT (**b**) at 26°C; Δ*rcsB* with WT (**c**), and $rcsD_{pe}$::$rcsD_{pstb}$ with WT (**d**) at 37°C.

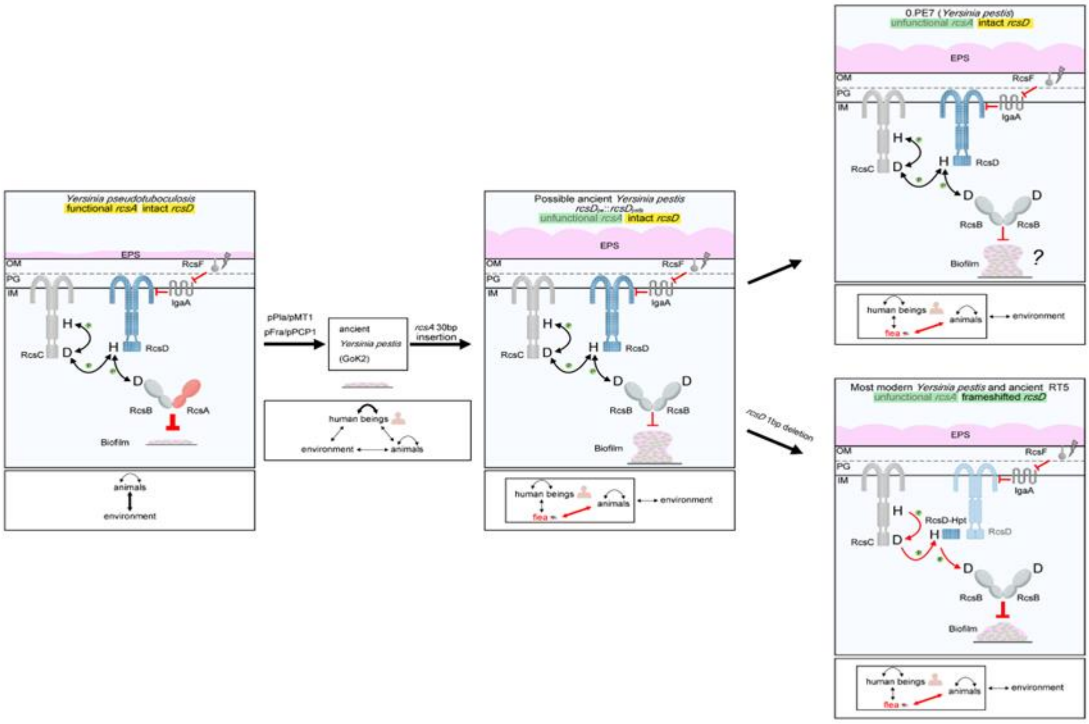


**Supplementary Figure 3. Evolutionary model of how genetic changes in the Rcs system fine tunes biofilm formation (upper panels) to preserve virulence of flea-bite transmitted (lower panels) *Y. pestis.***

Left: Functional *rcsA* and intact *rcsD* are encoded by ancestral *Y. pseudotuberculosis*. Biofilm formation was strongly repressed by the Rcs system due to the presence of RcsA. *Y. pseudotuberculosis* were transmitted between animals and the environment mainly by the fecal-oral route. Acquisition of pPla/pPCP1 and pFra/pMT1 plasmids led to the speciation of ancient *Y. pestis*, which were transmitted mainly between humans and occasionally by flea bites through early phase transmission. Once *rcsA* was pseudogenised, biofilm repression by *rcsA* was abolished, and flea-borne transmission was established. *rcsD* subsequently underwent a frameshift mutation in most modern strains, which reduced biofilm formation leading to low rate of spontaneous loss of *pgm* locus compared with an intermediate ancestral strain. pMT1 and pPCP1 refer to the two *Y. pestis*-specific plasmids that encode Ymt and Pla. In the bottom panels, arrows indicate transmission of bacteria between rodents, fleas and the external environment. Their weight indicates ecological importance.

OM: outer membrane; PG: peptidoglycan; IM: inner membrane; H: histidine; D:

aspartic acid; EPS: exopolysaccharides.

#### Supplementary Tables

**Supplementary Table 1. Strains used in this study.**

| **Strain** | **Genotype and description** | **Reference** |
|---|---|---|
| ***Yersinia pestis* KIM6+** | | |
| WT | Wild type of KIM6+ | Deng et al., 2002 |
| Δ*rcsD*$_{\text{N-term}}$ | Fragment between 1 and 1846 bp of *rcsD* deleted in KIM6+ | Sun et al., 2008 |
| *rcsD*$_{\text{pe}}$::*rcsD*$_{\text{pstb}}$ | *rcsD*$_{\text{pe}}$ substituted with *rcsD*$_{\text{pstb}}$ in KIM6+ | Sun et al., 2008 |
| Δ*rcsD* | Fragment between 1 and 2580 bp of *rcsD* deleted in KIM6+ | This study |
| Δ*rcsD*$_{\text{C-term}}$ | Fragment between 1898 and 2580 bp of *rcsD* deleted in KIM6+ | This study |
| Δ*rcsB* | *rcsB* gene deleted in KIM6+ | Sun et al., 2008 |
| *rcsD*$_{\text{pe}}$ (H844A) | *rcsD*$_{\text{pe}}$ mutated in KIM6+, at a site of conserved histindine of RcsD (RcsD$^{\text{H844A}}$) | This study |
| Δ*rcsD*$_{\text{N-term}}$ (H844A) | *rcsD*$_{\text{pe}}$ mutated in Δ*rcsD*$_{\text{N-term}}$, at a site of conserved histindine of RcsD (RcsD$^{\text{H844A}}$) | This study |
| *rcsD-hpt* (RBS*) | The predicted RBS of *rcsD-hpt* mutated in KIM6+ (C)AAAAGG > (A)ACAAAG) | This study |
| *rcsD* (ATT$^{312}$→GGT) | The predicted translation start codon of *rcsD-hpt* mutated in KIM6+ (ATT > GGT) | This study |
| *rcsD* (ATT$^{-312}$→ATG) | The predicted translation start codon of *rcsD-hpt* mutated in KIM6+ (ATT > ATG) | This study |
| Δ*rcsC* | *rcsC* gene deleted in KIM6+ | This study |
| *rcsD*$_{\text{pe}}$::*rcsD*$_{\text{pstb}}$ Δ*rcsC* | *rcsC* gene deleted in *rcsD*$_{\text{pe}}$::*rcsD*$_{\text{pstb}}$ | This study |
| Δ*rcsF* | *rcsF* gene deleted in KIM6+ | This study |
| *rcsD*$_{\text{pe}}$::*rcsD*$_{\text{pstb}}$ Δ*rcsF* | *rcsF* gene deleted in *rcsD*$_{\text{pe}}$::*rcsD*$_{\text{pstb}}$ | This study |
| Δ*igaA* | *igaA* gene deleted in KIM6+ | This study |
| *rcsD*$_{\text{pe}}$::*rcsD*$_{\text{pstb}}$ Δ*igaA* | *igaA* gene deleted in *rcsD*$_{\text{pe}}$::*rcsD*$_{\text{pstb}}$ | This study |
| Δ*rcsB* | *rcsB* gene deleted in KIM6+ | Sun et al., 2008 |
| *rcsB* (D56Q) | *rcsD*$_{\text{pe}}$ gene mutated in KIM6+, at a site of conserved aspartic acid of RcsB (RcsB$^{\text{D56Q}}$) | This study |
| ***Yersinia pestis* biovar Microtus strain 201** | | |
| WT | Wild type of biovar Microtus strain 201 | Song et al., 2004 |
| *rcsD*$_{\text{pe}}$::*rcsD*$_{\text{pstb}}$ | *rcsD*$_{\text{pe}}$ substituted with *rcsD*$_{\text{pstb}}$ in strain 201 | This study |
| Δ*rcsB* | *rcsB* gene deleted in strain 201 | This study |

**Supplementary Table 2. Plasmids used in this study.**

| Plasmid | Description | References |
|---|---|---|
| pUC19 (p)[a] | Cloning vector, ColE1 replicon , Amp$^r$ | Lab stock |
| $prcsD_{pe}$ | $rcsD_{pe}$ gene cloned in pUC19 | This study |
| $prcsD_{pstb}$ | $rcsD_{pstb}$ gene cloned in pUC19 | Sun et al., 2008 |
| $prcsD_{pe}$ (H844A) | $rcsD_{pe}$ gene with mutated conserved Histidine (H844A) cloned in pUC19 | This study |
| $prcsD_{C\text{-}term}$ | Fragment between 1828 and 2693 bp of $rcsD_{pe}$ cloned in pUC19 | This study |
| $prcsD_{C\text{-}term}$ (H844A) | The conserved histidine (H844A) mutated in $prcsD_{C\text{-}term}$ | This study |
| $prcsD_{C\text{-}term\ (648bp)}$, $prcsD_{2046\text{-}2693}$ | Fragment between 2045 and 2693 bp of $rcsD_{pe}$ cloned in pUC19 | This study |
| $prcsD_{pe}$ (TTG$^{-573}$$\rightarrow$CTT) | Point mutation at a site 573 bp upstream of stop codon in $prcsD_{pe}$ (TTG > CTT) | This study |
| $prcsD_{pe}$ (TTG$^{-462}$$\rightarrow$TTA) | Point mutation at a site 462 bp upstream of stop codon in $prcsD_{pe}$ (TTG > TTA) | This study |
| $prcsD_{pe}$ (TTG$^{-573}$$\rightarrow$CTT, TTG$^{-462}$$\rightarrow$TTA) | Points mutation at two sites 573 bp and 462 bp upstream of stop codon in $prcsD_{pe}$ (TTG > CTT, TTG > TTA) in $prcsD_{pe}$ | This study |
| p*rcsC* | *rcsC* gene cloned in pUC19 | This study |
| p*rcsC* (H489A) | *rcsC* gene with mutated HisKA domain (H489A) cloned in pUC19 | This study |
| p*rcsC* (D885A) | *rcsC* gene with mutated REC domain (D885A) cloned in pUC19 | This study |
| p*rcsC* (T913A) | *rcsC* gene with mutated REC domain (T913A) cloned in pUC19 | This study |
| p*rcsC* (D885A, T913A) | *rcsC* gene with mutated REC domain (D885A, T913A) cloned in pUC19 | This study |
| p*rcsC* (H489A, D885A) | *rcsC* gene with HisKA and REC domains (H489A, D885A) cloned in pUC19 | This study |
| p*rcsF* | *rcsF* gene cloned in pUC19 | This study |
| p*rcsF* (C125S) | *rcsF* gene with mutated domain (C125S) cloned in pUC19 | This study |
| p*igaA* | *igaA* gene cloned in pUC19 | This study |
| p*igaA* (C413S) | *igaA* gene with mutated periplasmic domain domain (C413S) cloned in pUC19 | This study |
| p*rcsB*-Flag | *rcsB* gene with fused C-terminal Flag tag cloned in pUC19 | This study |
| p*rcsB* (D56Q)-Flag | *rcsB* gene with mutated REC domain (D56Q) and fused C-terminal Flag tag cloned in pUC19 | This study |

| | | |
|---|---|---|
| p*rcsD*$_{pe}$-Flag | C-terminal Flag tag fused in p*rcsD*$_{pe}$ | This study |
| p*rcsD*$_{pe}$-Flag-His | C-terminal Flag and hexahistidine tags fused in p*rcsD*$_{pe}$ | This study |
| p*rcsD*$_{C\text{-}term}$-Flag-His | C-terminal Flag and hexahistidine tags fused in p*rcsD*$_{C\text{-}term}$ | This study |
| p*rcsD*$_{pstb}$-Flag | C-terminal Flag tag fused in p*rcsD*$_{pstb}$ | This study |
| pBAD/Myc-His A (pBAD) | Expression vector, Amp$^r$ | Invitrogen |
| pBAD*rcsD-hpt*$_{573bp}$ | Fragment between 2121 and 2693 bp of *rcsD*$_{pe}$ cloned in pBAD | This study |
| pBAD*rcsD-hpt*$_{462bp}$ | Fragment between 2230 and 2693 bp of *rcsD*$_{pe}$ cloned in pBAD | This study |
| pBAD/Myc-His A' (pBAD') | modified pBAD/Myc-His A without ATG | This study |
| pBAD'*rcsD-hpt* | Fragment between 2382 and 2693 bp of *rcsD*$_{pe}$ cloned in pBAD' | This study |
| pBAD' *rcsD-hpt* (H844A) | The conserved histidine (H844A) mutated in pBAD'*rcsD-hpt* | This study |
| pBAD' *rcsD-hpt* (ATT$^{-312}$→GGT) | The predicted translation start codon of *rcsD-hpt* mutated in pBAD'*rcsD-hpt* (ATT > GGT) | This study |
| *hmsT*::*lacZ* reporter | 350 bp of *hmsT* upstream sequence, together with the first seven codons of the *ORF* cloned in pGD926 | Guo et al., 2015 |
| pMal-*rcsD*$_{pstb}$-*lacZ* | The partial sequence of *rcsD*$_{pstb}$ (159 bp) containing the 8T (frameshifted region) cloned into pMAl-*lacZ* | This study |
| pMal-*rcsD*$_{pe}$-*lacZ* | The partial sequence of *rcsD*$_{pe}$ (158 bp) containing the 7T (frameshifted region) cloned into pMAl-*lacZ* | This study |
| pMal-*rcsD*$_{pe}$-stop-*lacZ* | An additional stop codon was introduced into the 158-bp *rcsD*$_{pe}$ sequence of pMal-*rcsD*$_{pe}$-*lacZ* | This study |
| pKD46 | λ red recombination vector for gene deletion | Datsenko et al., 2000 |
| pAC-crRNA-cm | vector for ligating crRNA and gene editing | Yan et al., 2017 |
| pKD46-Cpf1-amp | pKD46 with Cpf1, for gene editing | Yan et al., 2017 |

pUC19 (p)[a], represented as p for short. For p-derived plasmids, the fragment between 400 bp upstream of 5' UTR DNA region of target gene and 100 bp downstream of its 3' UTR DNA region was cloned into pUC19.

**Supplementary Table 3. Primers and oligos used in this study.**

| **Primers** | **Sequences** | **Function** |
|---|---|---|
| p*rcsC* F | GCTCTAGAGCAGAAGATGAACATGTTGTTGGT | Plasmid construction |
| p*rcsC* R | GGAATTCATTAGCCAAGTGGATAAAGAACTGT | |
| p*rcsD* F | CGGGATCCGCCGCCTAAACGGTGATCG | |
| p*rcsD* R | GGAATTCCGACAACGTTTACCCATTCAATT | |
| pBAD*rcsD-hpt*$_{573bp}$ F | CATGCCATGGTATTGTTAAATTCATTTGGTGCTAACT | |
| pBAD*rcsD-hpt*$_{573bp}$ R | GGAATTCTTATGGGTTACCTTCCTGCAACA | |
| pBAD*rcsD-hpt*$_{462bp}$ F | CATGCCATGGTATTGCTCGCGGCGGATG | |
| p*rcsD*$_{C\text{-}term}$ F | CGGGATCCGCGGGTATTTCTGATGAAGAGAT | |
| p*rcsD*$_{N\text{-}term}$ R | GGAATTCCTGGTTTTTAGTTGCCTCTCATAGAG | |
| *rcsC* deletion F | TCGGTTATCGGTAGTATATTCTGTTGTTAACACAGCCTGTCTGGCGATTCCGGGGATCCGTCGACC | Gene deletion by Recombineering |
| *rcsC* deletion R | ATTTTTGCCCCGCTCGACTCTCAAGCCCAGTTTTGCTCTGCAATACGTGTAGGCTGGAGCTGCTTCG | |
| *rcsD* deletion F | GCGCCAGAGCCGTCACGCGCTAACCAAGGGAATCGGCATTATCGTT ATTCCGGGGATCCGTCGACC | |
| *rcsD*$_{2580bp}$ down deletion R | CTTCAACCGATCGCCATCTGCGATATGCTGTTCTAATGTTTCACATAGTGTAGGCTGGAGCTGCTTCG | |
| *rcsD*$_{1898bp}$ up deletion F | TATGTAACCAATTATGTAAAAAATTAAACGGCCAATTAGAGATTC ATTCCGGGGATCCGTCGACC | |
| *rcsF* deletion F | TTGCTTCAATAGCACAACTTAAATGAATAAGGCTGAGGAATTTTATTCCGGGGATCCGTCGACC | |
| *rcsF* deletion R | TTAGATGAAACGTTCAGTGCTGAACCCTGACACACCGCTTGTTGA GTGTAGGCTGGAGCTGCTTCG | |
| *igaA* deletion F | TTTCATAGCGTTGACGTAAATTTAAAGCGCTCTGATGGGGATGGG ATTCCGGGGATCCGTCGACC | |
| *igaA* deletion R | CGTGCATCTGCTGGCATGAGCGATGATCAACCGGAGATGGGTGAGGTGTAGGCTGGAGCTGCTTCG | |
| *rcsD* fused 3xFlag F | GATTATAAAGATCATGACATCGACTACAAGGATGACGATGACAAGTGACCATGAACA | Overlap PCR for |

| | | |
|---|---|---|
| | ACCTTAATGTAATTATTGCTGATGAC | mutagenesis |
| *rcsD* fused 3xFlag R | GTCGATGTCATGATCTTTATAATCACCGTCATGGTCTTTGTAGTCTGGGTTACCTTCCTGCAACAGTCTG | |
| *rcsD* fused 3xFlag and His6 F | ATGACGATGACAAGCATCATCATCATCATCACTGACCATGAACA ACCTTAATGTAATTATTGCTG | |
| *rcsD* fused 3xFlag and His6 R | AGGTTGTTCATGGTCAGTGATGATGATGATGATGCTTGTCATCGTCATCCTTGTAGTCG | |
| *rcsC* (H489A) F | GCGGAATTGCGCACACCGCTGTATGGCATTATCGG | |
| *rcsC* (H489A) R | AGCGGTGTGCGCAATTCCGCACTAACGGTAGCCAGAAACATGGATTTTG | |
| *rcsC* (D885A) F | GCGGTGAACATGCCAAATATGGATGGTTACCGTTT | |
| *rcsC* (D885A) R | ATATTTGGCATGTTCACCGCGGTCAATACCATATCTACGGTATTAGTATTTAACGC | |
| *rcsC* (T913A) F | GCGGCGAATGCTTTGGCTGAGGGAAAACAAC | |
| *rcsC* (T913A) R | TCAGCCAAAGCATTCGCCGCTACACCAATAATCGGGAAATTGTGATTCAA | |
| *rcsD* (H844A) F | GCGCGATTAAAAGGCGTATTTGCCATGCTGAACC | |
| *rcsD* (H844A) R | AATACGCCTTTTAATCGCGCTACGGTTTGCGAAAGTGATGTTAAATCACTT | |
| *rcsB* (D56Q) F | AATTACTGAACTCTCTATGCCAGGGGATAAGTATGGTGATGGCATCAC | |
| *rcsB* (D56Q) R | AGAGTTCAGTAATTAGCACGTTGGCATCAAGTTTGGACAAATTGTTAATAAGC | |
| *rcsB* fused 3xFlag F | GATTATAAAGATCATGACATCGACTACAAGGATGACGATGACAAGTGAGCATGCCTGCAGGTCGACTCTAGA | |
| *rcsB* fused 3xFlag R | GTCGATGTCATGATCTTTATAATCACCGTCATGGTCTTTGTAGTCAAGCTTGGCGTAATCATGGTCATAGCTGTTTCCT | |
| *rcsD (*TTG$^{-462}$→TTA) F | TTACTCGCGGCGGATGAAACGGGGTTTCA | |
| *rcsD* (TTG$^{-462}$→TTA) R | GTTTCATCCGCCGCGAGTAATAATGTGTAATTATCATAATGTTGCGGATTATCCGTTATTGTA | |
| *rcsD (*TTG$^{-573}$→CTT) F | CTTCTAAATTCATTTGGTGCTAACTGCATTCTCACCGATGAAC | |
| *rcsD (*TTG$^{-573}$→CTT) R | GCACCAAATGAATTTAGAAGTATTGTAATGATCGAGCGTACT | |
| ig*aA* (C413S) F | CAGGGTAATGGCATGAGTTATGTCCCCCCCAACATTCAAAACACACG | |
| *igaA* (C413S) R | GTTGGGGGGGACATAACTCATGCCATTACCCTGAGCCTTCAGCATATCG | |

| | | |
|---|---|---|
| *rcsF* (C125S) F | TATCAACAAGCGGTGAGTCAGGGTTCAG CACTGAACGTTTCATCTAAATGA G | |
| *rcsF* (C125S) R | CAGTGCTGAACCCTGACTCACCGCTTGT TGATAGCAGCCAGGGAC | |
| Modified pBAD' F | TTTTTTGGGCTAACAGGAGGTTGGATCC CGGGGGTTCTCATCATCATCATCATCATG GTATGGC | Overlap PCR for ATG deletion in pBAD/Myc-His A |
| Modified pBAD' R | TGATGATGATGAGAACCCCCGGGATCCA ACCTCCTGTTAGCCCAAAAAACGGGTAT GGAGAAACA | |
| *rcsD* (H844) crRNA F | TGCAAACCGTACATCGATTAAAAGGCGT | Cas12a mediated genome editing |
| *rcsD* (H844) crRNA R | GCCTTTTAATCGATGTACGGTTTGCATC | |
| *rcsD* (H844A) oligo | TGATTTAACATCACTTTCGCAAACCGTAG CGCGATTAAAAGGCGTATTTGCCATGCT GA | |
| *rcsB* (D56) crRNA F | TCTGACCTCTCTATGCCAGGGGATAAGT | |
| *rcsB* (D56) crRNA R | TTATCCCCTGGCATAGAGAGGTCAGATC | |
| *rcsB* (D56Q) oligo | CAAACTTGATGCCAACGTGCTAATTACT CAGCTCTCTATGCCAGGGGATAAGTATG GTG | |
| *rcsD-hpt* (RBS)crRNA F | TTGTAATGTACTCAACCTTTTGTTGCGT | |
| *rcsD-hpt* (RBS)crRNA R | GCAACAAAAGGTTGAGTACATTACAATC | |
| *rcsD-hpt (*RBS*) Oligo | TTTATATCATCTTCTGTAATGTACTCAACT TTGTTTTGCGATTGTTCAGATAAAATTCG | |
| *rcsD*$_{\text{C-term}}$ (ATT$^{-312}$) crRNA F | TAG TACATTACAGAAGATGATATAAAGT | |
| *rcsD*$_{\text{C-term}}$ (ATT$^{-312}$) crRNA R | TTTATATCATCTTCTGTAATGTACTATC | |
| *rcsD*$_{\text{C-term}}$ (ATT$^{-312}$ →GGT) oligo | TGAACAATCGCAACAAAAGGTTGAGTA CGGTACAGAAGATGATATAAACCTCTATG A | |
| *rcsD* (ATT$^{-312}$→ATG) oligo | TGAACAATCGCAACAAAAGGTTGAGTA CATGACAGAAGATGATATAAACCTCTATG A | |
| *rcsD-lacZ* frameshift test F | ATGACCATGGCGGGTATTTCTGATGAAG | Construction of pMal-*rcsD*$_{\text{pe}}$ /*rcsD*$_{\text{pstb}}$-*lacZ* and pMal-*rcsD*$_{pe}$-stop-*lacZ* |
| *rcsD-lacZ* frameshift test F | GTACAGATCTTCTACTTTACTGCGAATCT CTAATTG | |
| *rcsD*$_{\text{pe}}$-stop-*lacZ* | GTACAGATCTTATGCTACTTTACTGCGAA TCTCTAATTG | |
| 16sRNA F | TGGGAGTGGGTTGCAAAAGA | qPCR |
| 16sRNA R | GTTACGACTTCACCCCAGTCA | |
| *ybtT* qPCR F | CGTTCAGAATGCAGATGCGG | |
| *ybtT* qPCR R | GAAGCATCCGTATCGCCTGA | |
| *osmY* qPCR F | ACGCTTTTCACGCCGTTTAC | |
| *osmY* qPCR R | GGTGTCGTACAGCTATCGGG | |
| *glnH* qPCR F | TCAGCAGTACGGTGTTGCAT | |
| *glnH* qPCR R | TTCAGCGTAAGTGCCGTCTT | |
| *tus* qPCR F | ACTGATGCTGACAGAACGCA | |

| | | |
|---|---|---|
| *tus* qPCR R | CGTGAGCAATGGAGGGAACT | |
| *hmsT* qPCR F | TGCTATCATCGTCGCCCAAG | |
| *hmsT* qPCR R | ATTTCTGCCCGTGTCGTTCT | |
| *igaA* qPCR F | CCATCCGGCGACGATTTTTC | |
| *igaA* qPCR R | CATTGTCACTCTGGGTCGCT | |
| *lsrF* qPCR F | ATTGGCATCCCACAGCAGAA | |
| *lsrF* qPCR R | TAGCCATGGTCAAAGGCCAG | |
| *irp2* qPCR F | TGGTAGCGATCTTCAGGGGA | |
| *irp2* qPCR R | ACCGACTTCTTCCAGCAAGG | |
| *pgm*-test F (*hmsS*) | CGCCCCTGATTTTTACGG | PCR for testing the loss of *pgm* locus |
| *pgm*-test R (*hmsS*) | CATCCCTGGCGTAAATGG | |
| *pgm*-test F (upstream) | CTTTAGATTGTATGTAGGCAAGAGA | |
| *pgm*-test R (downstream) | TGAGGCAGGTAGGTATCG | |

**Supplementary Table 4 (in an excel document). Differential expression of *Y. pestis* genes in *Y. pestis* mutant strains growing at temperatures (26°C or 37°C).**

Raw data for the RNA-seq libraries are stored in the National Center for Biotechnology Information Sequence Read Archive and have been assigned BioProject accession (PRJNA876755).

**Supplementary Table 5. Mutations in *Y. pestis* during evolution involved in this study.**

| Gene | ID | Mutation in *Y. pestis* | Effect | Occurrence |
|---|---|---|---|---|
| *rcsA* | YPTB2486 YPO2449 | 30-bp internal duplication in ORF | Derepression of HmsT caused by insertion of 10 residuals tandem repeat | All *Y. pestis*, except for branch 0 strain Pestoides A (0.PE4b) |
| *rcsD*[b] | YPO0919 | Indel (8T→7T) in *rcsD*C-term | Early termination of RcsD and accumulation of a small HPt encoded by *rcsD*C-term | Most modern strains, except for 620024, CMCC05009 (0.PE7), I-3086 and A-1825 |
| *rcsD*[c] | YPO0919 | Rearrangement, *rcsD*N-term and *rcsD*C-term -*rcsB*-*rcsC* were separated | Early termination of RcsD and accumulation of a small Hpt encoded by *rcsD*C-term | Nairobi (1. ANT); Anloga (0.PE3); Algeria3 (1.ORI) |
| *rcsD*[d] | YPO0919 | Indels in *rcsD*N-term | Early termination of RcsD and accumulation of a small HPt encoded by *rcsD*C-term | I-3086 (0.PE4m), A-1825 (2.MED1) |
| IS*ompC* | YPO1900, YPO1998 | Acquist of IS elements in *ompC* | Easy loss of *pgm* locus | Without IS: ancient, 91001, etc. (0.PE4;5; 2.MED);<br>With IS: CO92, KIM6+, etc |
| *igaA* | YPO0142 | Indels | Pseudogene | C-627, etc (0.PE4, 0.PE10, 0.PE5, 0.PE2, 2.MED) |
| *rcsC* | YPO0921 | Indel (8T→9T) | Pseudogene | Several strains (2.MED) |
| *rcsF* | YPO1070 | Indel | Pseudogene | D106004 (1.IN) |

*rcsD*[a], *rcsD*[b], and *rcsD*[c] stand for three different mutations of *rcsD* in *Y. pestis* during evolution as described in this Table.

**Supplementary Table 6. Putative SD and start codon sites in *rcsD* in Enterobacteriaceae.**

| Species | Predicted SD[e] | Spacer length between SD and start site | Predicted start site (fragment length of predicted *rcsD-hpt*) |
|---|---|---|---|
| *Escherichia coli* | AGGAAG | 14 bp | ATT (327 bp) |
| *Shigella boydii* | AGGAAG | 14 bp | ATT (327 bp) |
| *Shigella boydii* | GAGCAA | 4 bp | ATG (351 bp) |
| *Serratia fonticola* | AGAACA | 4 bp | ATG (357 bp) |
| *Proteus mirabilis* | GAGGC | 10 bp | TTG (372 bp) |
| *Yersinia frederiksenii* | AGGAAG | 7 bp | ATT (381 bp) |
| *Erwinia amylovora* | AGGACG | 8 bp | ATC (576 bp) |
| *Klebsiella quasipneumoniae* | AGGAGG | 14 bp | ATT (327 bp) |
| *Klebsiella quasipneumoniae* | AGCACG | 3 bp | ATG (351 bp) |

SD[e], Shine–Dalgarno sequence.
